# Regulation of Nuclear Transcription by Mitochondrial RNA

**DOI:** 10.1101/2022.12.10.519922

**Authors:** Kiran Sriram, Zhijie Qi, Dongqiang Yuan, Naseeb K. Malhi, Xuejing Liu, Riccardo Calandrelli, Yingjun Luo, Shengyan Jin, Ji Shi, Martha Salas, Runrui Dang, Brian Armstrong, Saul J. Priceman, Ping Wang, Jiayu Liao, Rama Natarajan, Sheng Zhong, Zhen B. Chen

**Affiliations:** Department of Diabetes Complications and Metabolism, Beckman Research Institute, City of Hope, 1500 Duarte Rd., Duarte, CA 91010, USA; Irell and Manella Graduate School of Biological Sciences, City of Hope, 1500 Duarte Rd., Duarte, CA 91010, USA; Department of Bioengineering, University of California San Diego, 9500 Gilman Dr., La Jolla, CA 92093, USA; Department of Genetics, Yale University School of Medicine, New Haven, CT 06510, USA; Translura, Inc. 5 Science park, New Haven, CT 06511, USA; Department of Stem Cell Biology and Regenerative Medicine, City of Hope, 1500 Duarte Rd., Duarte, CA 91010, USA; Department of Bioengineering, University of California Riverside, Riverside, CA 92521, USA; Departments of Hematology & Hematopoietic Cell Transplantation and Department of Immuno-Oncology, City of Hope, 1500 Duarte Rd., Duarte, CA 91010, USA; Department of Diabetes, Endocrinology & Metabolism, Beckman Research Institute, City of Hope, 1500 Duarte Rd., Duarte, CA 91010, USA

## Abstract

Chromatin-associated RNAs (caRNAs) form a relatively poorly recognized layer of the epigenome. The caRNAs reported to date are transcribed from the nuclear genome. Here, leveraging a recently developed assay for detection of caRNAs and their genomic association, we report that mitochondrial RNAs (mtRNAs) are attached to the nuclear genome and constitute a subset of caRNA, which we termed mt-caRNA. In four human cell types analyzed, mt-caRNAs preferentially attach to promoter regions. In human endothelial cells (ECs), the level of mt-caRNA-promoter attachment changes in response to environmental stress that mimics diabetes. Suppression of a non-coding mt-caRNA in ECs attenuates stress-induced nascent RNA transcription from the nuclear genome, including that of critical genes regulating cell adhesion, and abolishes stress-induced monocyte adhesion, a hallmark of dysfunctional ECs. Finally, we report increased nuclear localization of multiple mtRNAs in the ECs of human diabetic donors, suggesting many mtRNA translocate to the nucleus in a cell stress and disease-dependent manner. These data nominate mt-caRNAs as messenger molecules responsible for mitochondrial-nuclear communication and connect the immediate product of mitochondrial transcription with the transcriptional regulation of the nuclear genome.

## Introduction

Mitochondria are key cellular organelles that control energy production, stress response, and cell fate. Mitochondrial fitness is paramount to cellular homeostasis and organismal health, and its dysfunction is a common denominator of numerous diseases ranging from metabolic, cardiovascular, to neurological disorders (*1*). As derivatives of endosymbiotic bacterial ancestors, mitochondria in eukaryotes possess their own circular and double-stranded genome (mtDNA) of 16.5 kb in size. mtDNA is bidirectionally transcribed to produce 13 mRNAs, 2 rRNAs, and 22 tRNAs involved in oxidative phosphorylation (OXPHOS). The mitochondrial genome also produces non-coding RNAs (ncRNAs) including small RNAs and long ncRNAs (lncRNAs), whose functions are yet to be fully characterized (*2–4*).

As a critical hub for cellular processes and signaling pathways, mitochondria coordinate with the nucleus to regulate gene expression and cell function (*5*). On the one hand, the nucleus communicates to the mitochondria through anterograde signaling. For example, nucleus-encoded transcription factors (TFs) such as NRF1 and TFAM, co-regulators such as PGC1α, and the RNA polymerase POLRMT regulate mitochondrial biogenesis, mtDNA replication and transcription (*6–10*). Nucleus-transcribed non-coding RNAs (ncRNAs) have also been shown to regulate mitochondrial RNA processing and translation (*11–13*). Conversely, mitochondria can communicate with nucleus through mitochondrial retrograde signaling, typically mediated by reactive oxygen species, ATP, and Ca^2+^ (*14–16*). Mitochondria-derived short peptides have also been shown to regulate nuclear gene expression (*17*). Moreover, mitochondria DNA (mtDNA) and double-stranded RNAs (dsRNAs) can also be released into the cytoplasm, which turn on an endogenous “danger signal” to trigger a type-I interferon response and innate immune surveillance (*18–22*). However, it is unclear whether the immediate output of mitochondrial transcription, i.e., the mitochondrial RNAs (mtRNAs), can act as retrograde signaling messengers to communicate directly with the nucleus.

Several mtRNAs have been detected in the nucleus. Burzio et al detected *MT-RNR2-* derived sense noncoding mitochondrial RNA (SncmtRNA) in the nucleus of human endothelial cells (ECs) and several cancer cell lines (*23–25*). Perry et al identified a small mitochondrial-encoded ncRNA, mito-ncR-805, localized in the nucleus of alveolar epithelial cells, and correlated with the increased expression of nuclear-encoded genes that regulate mitochondrial function (*26*). Yet to be clarified is whether any nuclear-translocated mtRNA can alter the epigenome, and subsequently impact gene transcription and cellular behavior.

The chromatin-associated RNAs (caRNAs) form a critical layer of the epigenome that can regulate nuclear transcription (*27, 28*). Here, we show that caRNAs are not exclusively nuclear genome-produced RNA – they also include mtRNA. The chromatin-associated mtRNAs (mt-caRNAs) preferentially attach to the promoter regions of the nuclear genome. In human ECs, the mtRNA-chromatin attachment levels change in response to cellular stress induced by high glucose and TNFα (HT). As an example, we identify the mitochondrial lncRNA SncmtRNA as a mt-caRNA. In human ECs, suppression of SncmtRNA attenuates HT stress induction of nascent RNAs transcribed from the nuclear genome, including the cell adhesion molecules ICAM1 and VCAM1, and abolishes stress-induced monocyte-EC adhesion. In addition to SncmtRNA, we show nuclear localization of other mtRNAs including MT-CYB and MT-ND5 in human ECs, which is increased by HT and in diabetic human donors. Collectively, our findings suggest that mtRNAs are a class of retrograde messengers that mediates cellular response by attaching to chromatin and regulating nuclear transcription.

## Results

### Pervasive association of mtRNA with chromatin

To test whether mtRNAs are attached to the nuclear genome, we generated high-resolution RNA-genome interaction maps from human embryonic stem cells (H1), foreskin fibroblasts (HFF), and lymphoblasts (K562) (*29*) using our recently developed iMARGI (in situ mapping of RNA-Genome Interactions) technology (*30, 31*). By simultaneously sequencing caRNAs and their associated genomic DNA sequences from purified nuclei (*32*), iMARGI can differentiate the sequencing reads originating from RNA (RNA-end reads) and genomic DNA (DNA-end reads) in the chimeric reads. As expected, iMARGI data revealed diverse caRNAs transcribed from thousands of coding and non-coding genes as well as transposable elements in the nuclear genome (*29*).

Surprisingly, in each cell type profiled, more than 6% of iMARGI-mapped read pairs had the RNA-end reads uniquely mapped to mtRNAs. These large numbers of mtRNA reads cannot be due to errors mapping to the nuclear DNA of mitochondrial origin (NUMT) (*33*), because 98.7% of mtRNA-matching sequences contain more than 5 bp mismatches to any NUMTs. Based on the normalized RNA-end reads, mtRNAs exhibited higher chromatin association level (CAL) than that of nuclear genome-transcribed RNAs (nuRNAs) in all three cell types (p = 0.0022, t test) (**Fig. 1A**). iMARGI also revealed the attachment of mtRNAs to various genomic regions, including promoters, coding sequences (CDS), 5’ and 3’ untranslated regions (UTRs), (super-)enhancers, introns, and intergenic regions, with promoters showing the highest enrichment of mtRNA association in all three cell types (largest p = 0.0074, one-way ANOVA) (**Fig. 1B-D**). SncmtRNA (GenBank: DQ386868.1), a mitochondria-encoded lncRNA that has been observed to be nuclear localized (*23, 24*), is also one of the caRNAs and showed higher CALs than that of nuRNA (p-value = 0.00017, student’s t-test) (**Fig. 1A**) and enriched association with promoters as compared to other genome regions (**Fig. 1E-G**).

**Figure 1.**
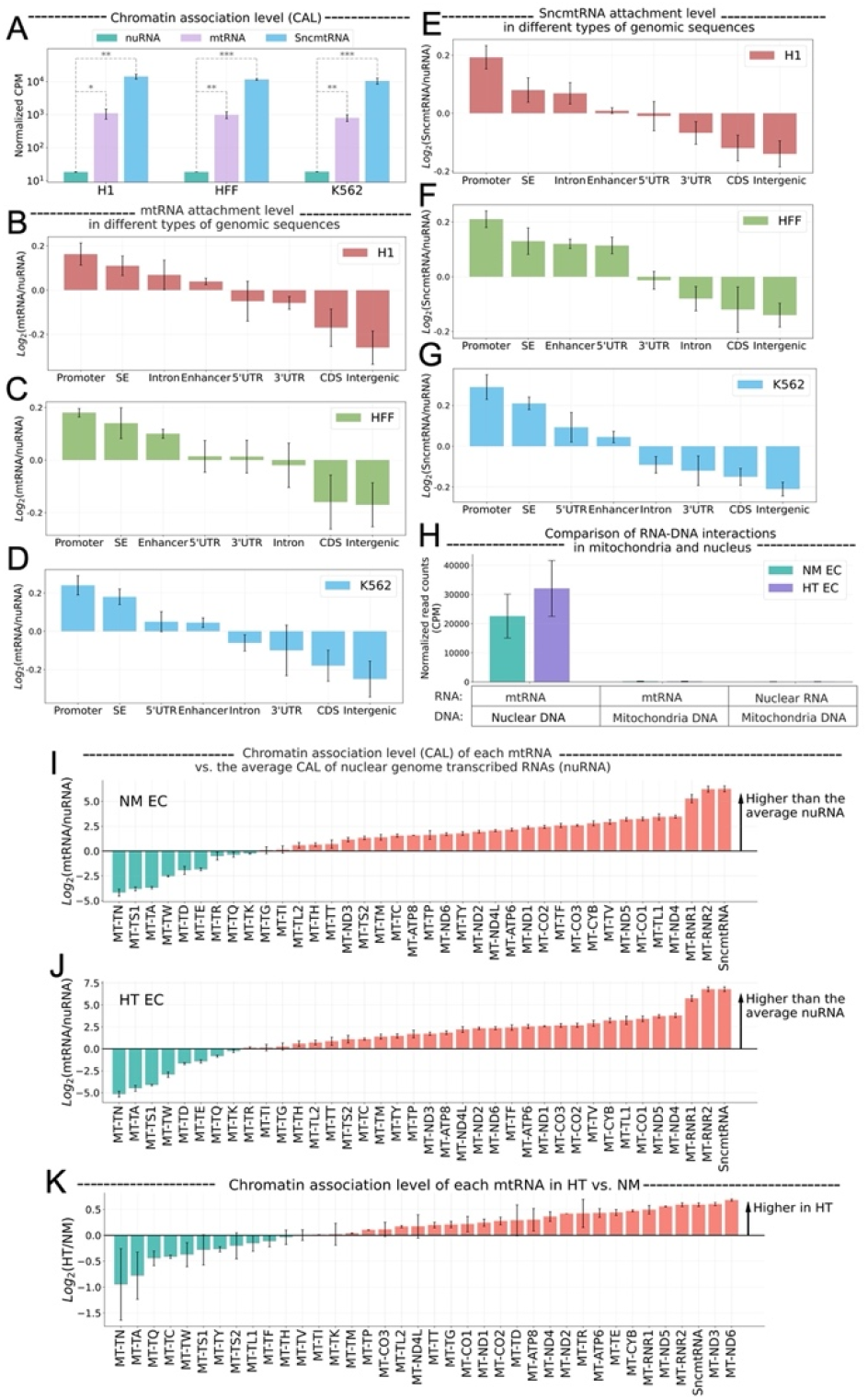
Association of mtRNA with chromatin. **(A)** Chromatin association levels (CAL) of nuclear genome transcribed RNA (nuRNA) (blue), mtRNA (green), and the SncmtRNA (purple) in H1 embryonic stem cells (H1), foreskin fibroblasts (HFF), and K562 lymphoblasts (K562) (columns). CAL is estimated by the normalized Counts Per Million (CPM) of iMARGI RNA-end read counts. **: p-value < 0.01, ***: p-value < 0.001. **(B-D)** Normalized CAL of mtRNA (y axis) on 8 types of genomic sequences (columns) in H1 (B), HFF (C), and K562 (D). Log2(mtRNA/nuRNA): the log ratio of mtRNA and nuRNA read counts in the iMARGI RNA-end reads. SE: super enhancer. UTR: untranslated region. CDS: coding sequence. **(E-G)** Normalized CAL of the SncmtRNA (y axis) on 8 types of genomic sequences (columns) in H1 (E), HFF (F), and K562 (G). Log2(SncmtRNA/nuRNA): the log ratio of SncmtRNA and nuRNA read counts in the iMARGI RNA-end reads. **(H)** Normalized numbers of iMARGI read pairs (y axis) mapped to mtRNA and nuclear DNA (first two columns), mtRNA and mitochondrial DNA (middle two columns), and nuRNA and mitochondrial DNA (last two columns) in normal mannitol control (NM)-(green) and high glucose and TNFa (HT)-treated (purple) endothelial cells (ECs). **(I-J)** Comparison of the CAL of each mtRNA (column) and the average CAL of all nuRNA in NM (I) and HT EC (J). Y axis: the log ratio of mtRNA and nuRNA read counts in the iMARGI RNA-end reads. y > 0 indicates that this mtRNA (column) exhibits a higher level of chromatin association than the average CAL of the nuRNA. **(K)** Comparison of the CAL of each mtRNA (column) between NM- and HT-treated EC. Log2(HT/NM): log ratio of CAL between HT and NM per each mtRNA. y >0 indicates higher CAL in HT vs NM.

Next, we investigated whether the chromatin-mtRNA association changes under a cellular stress condition. To this end, we analyzed iMARGI data from human umbilical vein ECs (HUVECs) under normal glucose (NM) and combined treatment of high glucose and TNFα (HT), which we have used to mimic healthy vs diabetic conditions (*34*). While ECs under NM stressed control show comparable levels of chromatin-attached mtRNAs as the other three cell types, the CALs of mtRNA increased from NM to HT (p-value = 0.037, student’s t test) (**Fig. 1H**). In contrast, under both conditions, read pairs between nuRNA and mitochondrial DNA (mtDNA) or mtRNA and mtDNA were barely detected (**Fig. 1H**), suggesting minimal mitochondrial contamination in the iMARGI libraries.

Quantification of the CAL of mtRNAs, including coding and non-coding transcripts, indicated that the majority of mtRNAs exhibit not only abundant but also higher levels of chromatin association than the nuRNAs under both conditions in ECs (**Fig. 1I,J**). Moreover, comparison between NM vs HT revealed that HT further induced the CALs of most of the mtRNAs (**Fig. 1K**). Specifically, SncmtRNA exhibited a 64-fold higher CAL than the average CAL of all nuRNAs in NM (p = 0.00082, student’s t test) (**Fig. 1I**), which was further increased by HT (p = 0.034, student’s t test) (**Fig. 1K**). Taken together, these data identify mtRNA as a component of ca<u>RNA</u> (hereafter called <u>mt-caRNA</u>) with preferential genomic association with promoters in multiple cell types. Furthermore, the mtRNA-chromatin association can be affected by a stress condition, exemplified by ECs under HT and numerous mtRNAs including SncmtRNA.

### Effect of SncmtRNA perturbation on the expression and function of nuclear-encoded genes in ECs

The chromatin association of mtRNAs, especially with promoters, prompted us to posit the involvement of mt-caRNAs in nuclear transcription. To test this hypothesis, we elected SncmtRNA in ECs as an example, considering that 1) SncmtRNA show significantly enriched CAL in ECs, which is further increased by HT in the iMARGI data (**Fig. 1I-K**) and 2) perturbation of a lncRNA is less likely to directly impact mitochondrial function than targeting a mitochondria-encoded mRNA, rRNA, or tRNA.

We first confirmed the presence and chromatin association of SncmtRNA in ECs. As described (*23, 24*), SncmtRNA is comprised of 1559 nt sequence encoding mitochondrial 16S rRNA (transcribed from the H strand of mitochondrial genome), with its 5’ end appended to an 815 nt invert repeat, forming a putative double-stranded hairpin-shaped structure (**Fig. 2A, Fig. S1A**). Based on this description, we verified the presence of SncmtRNA in ECs using reverse transcription (RT)-PCR with primers flanking the putative hairpin loop and the unique junction region (**Fig. 2A,B**), followed by Sanger sequencing (**Fig. S1B**). As a control, PCR without RT failed to amplify SncmtRNA, evident of an RNA. Moreover, in Rho^0^ cells that are depleted of mtDNA (*35*), SncmtRNA, like 16S rRNA, was not detected (**Fig. 2B**), indicative of its mitochondrial origin.

**Figure 2.**
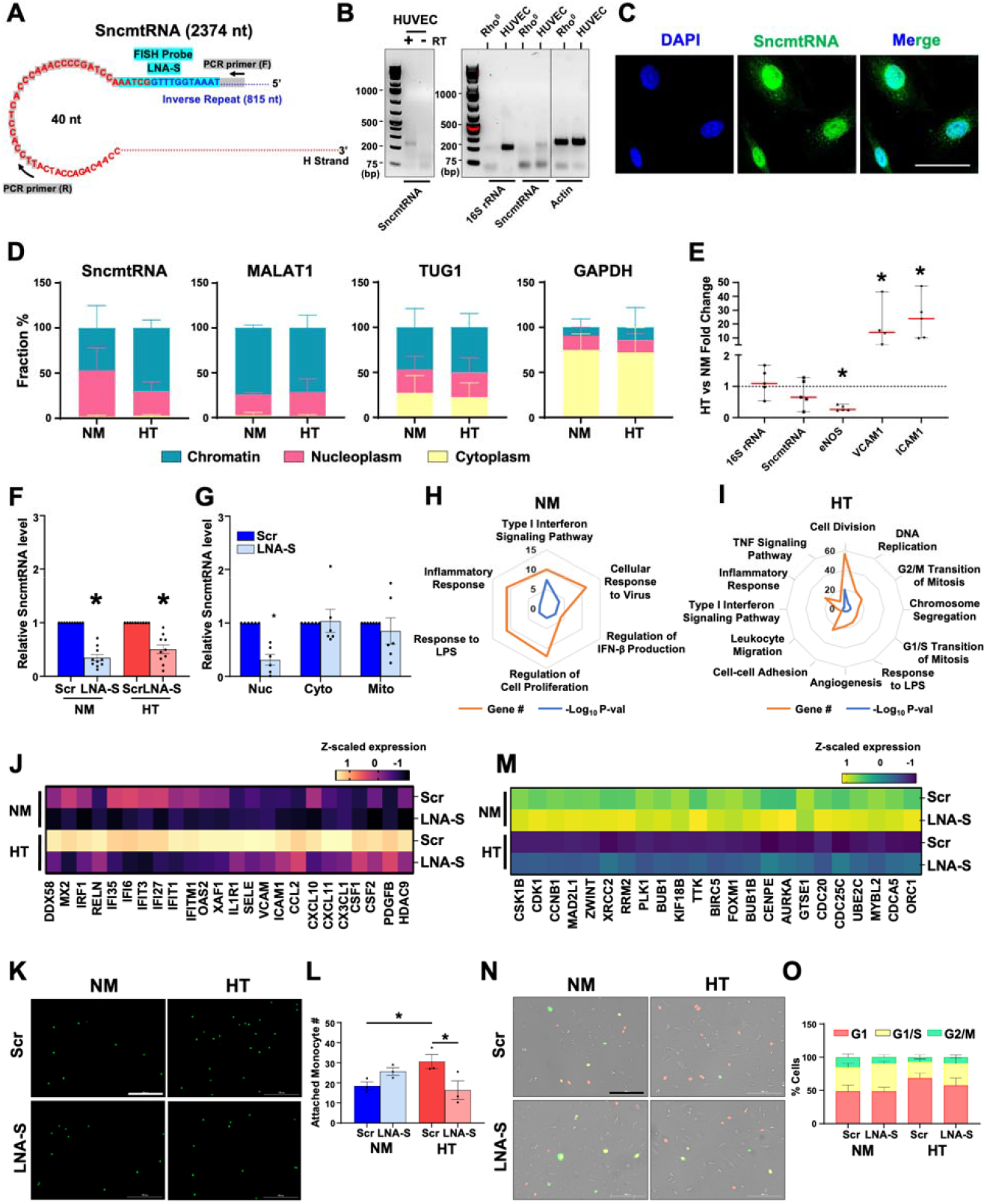
SncmtRNA knockdown affects EC expression and function. (**A**) Putative SncmtRNA structure and the location of LNA-GapmeR target region (LNA-S) and PCR primers. (**B**) PCR of SncmtRNA. Left: ECs with (+) or without (-) reverse transcription (RT). Right: ECs in comparison with Rho^0^ cells, with 16S rRNA and β-actin detected as controls. (**C**) FISH images of SncmtRNA in ECs, with DAPI staining nuclei. Scale bar = 50 μm. (**D, E**) ECs were maintained in normal glucose + 25 mM Mannitol (NM) or high-glucose + TNFα (HT) for 72 hr. (D) RT-qPCR of respective RNAs in subcellular fractions. Data plotted as mean ± SEM from three independent experiments. (**E**) RT-qPCR of various RNAs in whole cells, with levels in NM set at 1. Data represent mean ± SEM from five independent experiments. (**F,G**) RT-qPCR of SncmtRNA in (F) total ECs transfected with scramble (Scr) or SncmtRNA-targeting (LNA-S) LNA GapmeRs under NM and HT and in (G) subcellular fractions (nucleus, cytoplasm, and mitochondria) from ECs transfected with Scr or LNA-S and kept under NM. Data plotted as mean ± SEM from six independent experiments. (**H-O**) ECs treated as in (F) were profiled by bulk RNA-seq. **(H,I)** Gene Ontology (GO) terms enriched in DEGs due to Snc-KD under NM or HT. (**J,M**) Heatmap showing z-scaled expression in transcripts per million (TPM) of innate immune and inflammatory genes induced by HT and suppressed by Snc-KD in (J) and cell cycle genes suppressed by HT and rescued upon Snc-KD in (M). (**K,L,N,O**) Representative images and quantification of fluorescently labeled monocytes adhesion to HUVECs (in K,L) and FUCCI assay showing ECs in various phases of cell cycle (in N,O). Scale bar = 200 μm. Data plotted as mean □±□SEM from three and five independent experiments. *P<0.05 based on Student’s t test as compared to NM (in E) or Scr (in F,G,L).

We also performed fluorescence *in-situ* hybridization (FISH) with an antisense locked nucleic acid (LNA) probe targeting the unique junction region of SncmtRNA, which detected abundant signals in the nucleus in ECs. FISH also detected SncmtRNA signals in the cytoplasm resembling the morphology of mitochondria (**Fig. 2C**). As a FISH control, probe with scrambled sequence did not reveal specific signals (**Fig. S1C**). Consistently, qPCR of SncmtRNA in different EC subcellular fractions detected the majority of SncmtRNA in the nuclear fractions including chromatin and nucleoplasm, with MALAT1 (predominantly nuclear localized), GAPDH (predominantly cytoplasm-localized), and TUG1 (localized in both nucleus and cytoplasm) quantified as controls (**Fig. 2D**). Moreover, compared to NM, HT increased chromatin-associated SncmtRNA in ECs, while there was no change for MALAT1, TUG1, and GAPDH (**Fig. 2D**).

The HT-increased mt-caRNA, including SncmtRNA could be due to the increased total levels of mtRNAs in ECs. Arguing against this possibility, high glucose and TNFα are known to impair mitochondrial biogenesis (*36, 37*). Consistently, bulk RNA-seq of ECs under NM vs HT revealed that several nuclear-encoded genes regulating mitochondrial biogenesis, such as Nuclear Respiratory Factor 1 (*NRF1*) and Transcription Factor A, Mitochondrial (*TFAM*) were downregulated (**Fig. S1D**). Additionally, mitochondria visualized by Mitotracker showed fragmentation under HT (**Fig. S1E**). In terms of mitochondria-encoded transcripts, the majority remained unchanged (**Fig. S1D**). Similarly, the total RNA levels of SncmtRNA as well as 16S rRNA and antisense mitochondrial non-coding RNA (ASncmtRNA), all derived from *MT-RNR2*, were comparable between NM and HT. In contrast, the expression of eNOS, hallmark of EC homeostasis was decreased, and that of VCAM1 and ICAM1, typical inflammation markers were strongly induced by HT (**Fig. 2E**). Therefore, the pervasive increase of mt-caRNAs is unlikely due to a global increase of total mtRNAs.

Upon confirming the presence, sequence, and subcellular localization of SncmtRNA, we designed three SncmtRNA-targeting LNA GapmeRs, which can cause RNase H-mediated RNA degradation in the nucleus (*38*). While LNA-S1 targets the unique chimeric junction in SncmtRNA, LNA-S2 and S3 target different regions of H strand portion (**Fig. S2A**). As expected, only LNA-S1 (hereafter termed LNA-S) consistently decreased SncmtRNA level without affecting those of 16S rRNA and ASncmtRNAs (**Fig. S2B**) and was thus used in the subsequent SncmtRNA knockdown (Snc-KD) experiments. While LNA-S decreased the total cellular SncmtRNA in ECs under both NM and HT (**Fig. 2F**), the inhibitory effect was primarily in the nucleus, as LNA-S had insignificant impact on cytoplasmic and mitochondrial SncmtRNA (**Fig. 2G**). Therefore, any consequence observed with LNA-S should be primarily due to the reduction of SncmtRNA in the nucleus, including chromatin associated-SncmtRNA.

Bulk RNA-seq revealed that under NM, Snc-KD resulted in 265 differentially expressed genes (DEGs), including 160 down- and 105-upregulated (**Fig. S2C**). Under HT, Snc-KD led to more DEGs, i.e. 732 DEGs (403 down- and 329 up-regulated) in ECs (**Fig. S2D**), with 120 common DEGs between NM and HT (**Fig. S2E**). Pathway enrichment analysis indicated that innate immune and inflammatory responses are the most consistently affected pathways under both NM and HT, with more pathways involved in innate immune response under NM (e.g. Type I Interferon signaling and cellular response to virus and LPS) and more pathways involved in inflammatory response under HT (e.g., TNF signaling, leukocyte migration, and cell-cell adhesion) (**Fig. 2H,I**). Moreover, these pathways are mainly enriched in the downregulated DEGs by Snc-KD (**Fig. S2G,I**). For example, hallmark genes for innate immune and inflammatory activation, including *DDX58, MX2, VCAM1*, and *ICAM1*, were strongly suppressed by Snc-KD under both NM and HT (**Fig. 2J**).

Consistent with the gene expression changes, the HT-induced pro-inflammatory response of ECs assayed by monocyte adhesion, was abrogated by Snc-KD (**Fig. 2K,L**). In contrast, cell proliferation, cell division, and cell cycle regulation were mostly upregulated by Snc-KD, especially under HT (**Fig. 2I, Fig. S2H**), evident by the significant induction of cell cycle regulators, e.g., *CDK1* and *CCNB1* (**Fig. 2M**). FUCCI assay, which detects cell cycle progression and division (*39*), revealed while HT caused a cell cycle arrest at G1 phase, LNA-S led to a trend towards promoting the G1-arrested cells to S phase (**Fig. 2N,O**). In contrast to the profound effect of Snc-KD in nucleus-encoded transcriptome, there was no significant change in mtRNA levels caused by Snc-KD. Neither did Snc-KD have a clear effect on mitochondria content or morphology (**Fig. S2J**). Taken together, these data suggest that the mitochondria-encoded and partially chromatin-associated SncmtRNA plays a role in the regulation of nuclear-encoded transcripts.

### SncmtRNA regulates transcription of the nuclear genome

To further test our hypothesis that SncmtRNA regulates the transcription of nuclear-encoded genes, we reasoned that the effect of Snc-KD should be reflected by changes in the levels of nascent RNA, the immediate products of transcription. Furthermore, SncmtRNA transcriptional targets should show consistent changes between the nascent RNA levels in the nuclei and the total RNA levels in whole cells by Snc-KD. Thus, we performed single-nuclear and single cell RNA-seq (snRNA and scRNA-seq) in ECs with or without Snc-KD and subjected to HT to capture simultaneously the nuclear and the whole cell transcriptome at single cell level. While scRNA-seq generated the majority of reads mapped to exons, snRNA-seq generated 40–50% of reads mapped to introns (**Fig. S3A,B**), enabling better interrogation of nascent RNAs. On average, 5500 cells/sample were sequenced for snRNA-seq and 3500 cells/sample were sequenced for scRNA-seq, both of which performed with biological replicates. While snRNA- and scRNA-seq were highly correlated (Spearman correlation coefficient [SCC] = 0.9) (**Fig. 3A-C** and **Fig. S3C,D**), snRNA-seq identified 462 DEGs (comprising 302 down- and 160 up-regulated) and scRNA-seq returned 318 DEGs (comprising 174 down- and 144 up-regulated) as a result of Snc-KD in HT (**Fig. S3E,F**). Considering the comparable average reads per cell and total genes detected by sc- and snRNA-seq, but substantially lower median UMI counts per cell detected by snRNA-seq (**Fig. S3G,H**), the nuclear transcriptome appears to show a greater change than the whole-cell transcriptome due to Snc-KO, supporting a role of SncmtRNA in nuclear transcription.

**Figure 3.**
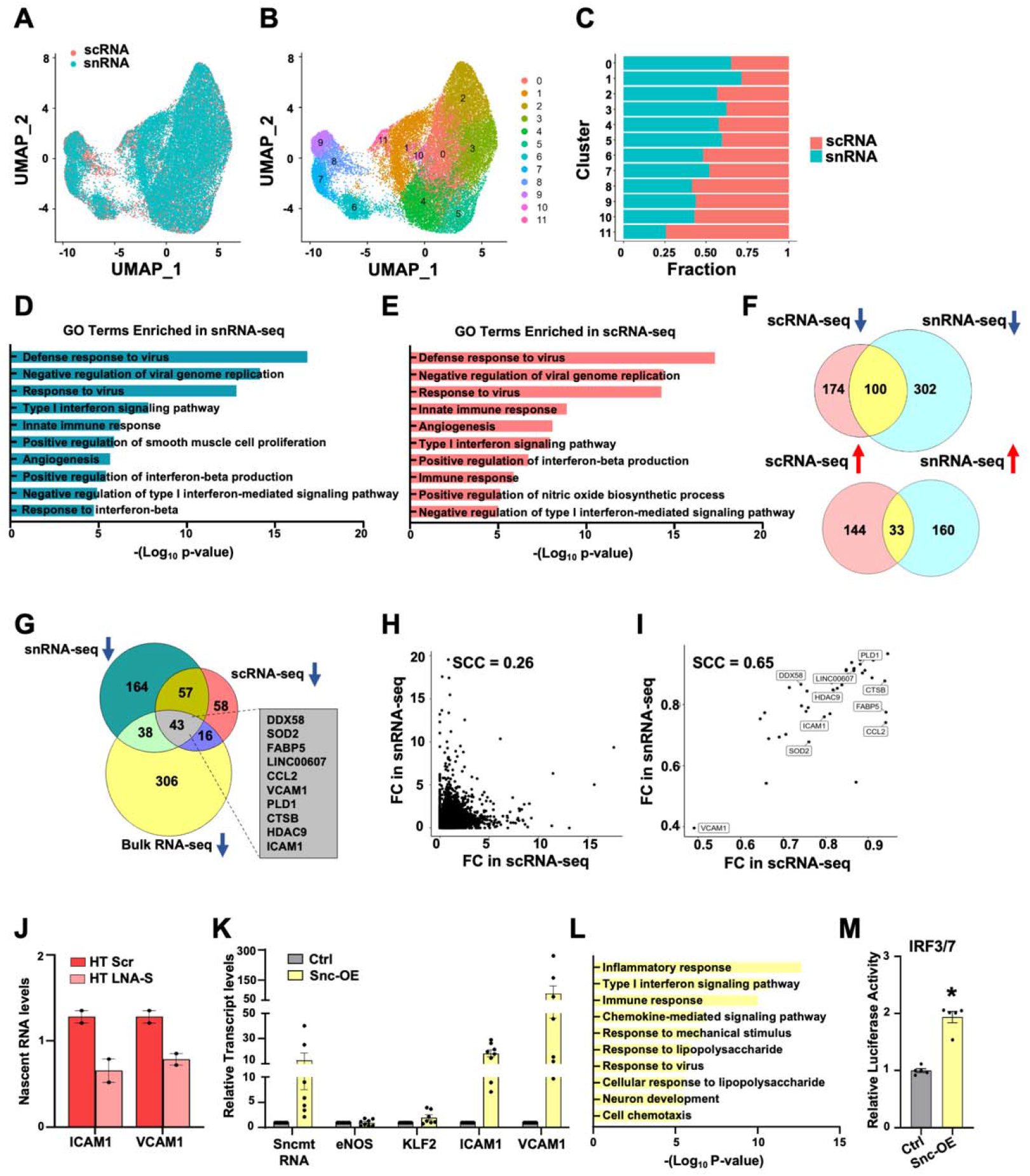
SncmtRNA regulates the transcription of nuclear genes. (**A-J**) ECs were transfected with Scr or LNA-S and exposed to HT for 72 hrs in biological replicates and then subjected to scRNA and snRNA-seq. UMAP embedding colored by cells (scRNA) or nuclei (snRNA) in (**A**) or by unsupervised clustering (**B**). (**C**) Fraction of cells (scRNA) or nuclei (snRNA) in each cluster from (B). (**D, E**) GO terms enriched by Snc-KD in snRNA-seq (**D**) or scRNA-seq (**E**). (**F**) Venn diagrams showing the number of common down or up-regulated DEGs between sc- and snRNA-seq. (**G**) Venn diagram showing the number of DEGs commonly down-regulated by Snc-KD in three RNA-seq datasets. Top ranked DEGs involved in innate immune and inflammatory response among the 43 intersecting genes are shown. (**H, I**) Correlation plot between Snc-KD-caused fold change (FC) in sn- and scRNA-seq of the commonly detected genes excluding the 43 intersecting genes (H) or the 43 intersecting genes (**I**). SCC are indicated. (**J**) Nascent RNA levels of ICAM1 and VCAM1 were quantified by RT-qPCR in the same batches of ECs used for sc- and snRNA-seq. (**K,L**) ECs were infected with control AAV (Ctrl) or AAV-driven SncmtRNA overexpression (Snc-OE) for 48 hrs. (**K**) RT-qPCR of respective transcripts. (**L**) Enriched pathways (GO terms) by Snc-OE identified from bulk RNA-seq. (**M**) Quantification of the activity of luciferase reporter driven by promoter containing IRF3/7 binding sites upon 293T cells infected by AAV (Ctrl) or Snc-OE.

Next, we identified the potential transcriptional targets of SncmtRNA by analyzing DEGs consistent between snRNA- and scRNA-seq datasets. Among common DEGs shared by sc- and snRNA-seq, 100 were downregulated and 33 were upregulated by Snc-KD, suggesting that SncmtRNA is more likely to be a transcriptional activator than suppressor (**Fig. 3F**). Among these common DEGs, 43 were consistently downregulated in bulk RNA-seq and sn- and scRNA-seq, including *ICAM1, VCAM1, CCL2*, and *DDX58* (**Fig. 3G, Fig. S4A**). These genes showed higher correlation between snRNA and scRNA-seq (SCC=0.65) as compared to other DEGs (SCC=0.26) (**Fig. 3H,I**), suggesting that these genes are more likely to be regulated at the transcriptional level by SncmtRNA. Interestingly, many of these genes are involved in innate immune and inflammatory response, which were the most enriched pathways shared by DEGs identified from snRNA- and scRNA-seq (**Fig. 3D,E**). In contrast, 29 genes were consistently upregulated upon Snc-KD in the bulk RNA-seq and scRNA-seq but not in the snRNA-seq data (**Fig. S4B,C)**. These genes may also be regulated by SncmtRNA but not via transcriptional regulation. Indeed, KLF2 is one such gene and its expression appears primarily regulated post-transcriptionally through miRNA-92a, which occurs in the cytoplasm (*40, 41*).

To test our hypothesis, we performed nascent RNA pulldown assay to demonstrate that Snc-KD indeed led to a reduction of nascent transcripts of ICAM1 and VCAM1 in ECs exposed to HT (**Fig. 3J**). Complementarily, we overexpressed SncmtRNA (Snc-OE) in ECs using AAV9, which enters the nucleus and drives the transcription of SncmtRNA (*42*). Snc-OE strongly induced ICAM1 and VCAM1, but not that of eNOS and KLF2 (**Fig. 3K**). At a transcriptome level, Snc-OE resulted in 553 DEGs (289 down- and 264 up-regulated), among which similar pathways to those affected by Snc-KD, e.g. Type I interferon signaling and responses to virus and lipopolysaccharide were also enriched (**Fig. 3L**). Consistently, Snc-OE in HEK-293 cells also increased the luciferase activity driven by promoters containing binding elements of IRF3/7, key TF mediating Type-I IFN signaling (**Fig. 3M**). Collectively, these data suggest that SncmtRNA regulates the transcription of nuclear-encoded genes, exemplified by those promoting innate immune and inflammatory response.

### Nuclear localization of mitochondrial transcripts

To validate our findings on mt-caRNA beyond SncmtRNA, we extended our study to other mt-caRNAs identified from iMARGI analysis. Specifically, we performed single molecule RNA FISH (smFISH) in ECs for MT-CYB and MT-ND5, which showed high CAL (**Fig. 1, I-K**) that was further increased by HT in ECs and have sequences feasible for smFISH probe design. We first used RNAscope, which is based on target-specific double Z probes (*43*) designed against MT-CYB and MT-ND5 respectively. We ensured that the sequences used for design of RNAscope probes did not match any nuclear genome at 100%, minimizing the possibility of detecting signals from NUMTs. While the positive control (*POLR2A*, a universally expressed gene) detected strong punctate signals and the negative control (*dapB*, a bacterial gene not expressed in humans) yielded no specific signals (**Fig. S5A,B**), smFISH for MT-CYB and MT-ND5 detected signals largely overlapping with that of Mitotracker (**Fig. S5C,D**). In contrast, smFISH with Rho^0^ cells showed no signals of either of the mtRNAs, indicating that the probes indeed detected mitochondrial transcripts (**Fig. S5E**).

At higher magnification, confocal microscopy revealed a fraction of MT-CYB and MT-ND5 signals in the nuclei as clear puncta, which overlapped with that of DAPI in ECs under NM and significantly increased by HT (**Fig. 4A-C**). Three-dimensional reconstruction of the confocal images visualized the MT-CYB signals embedded in the DAPI-stained nuclei in ECs under NM, which became more pervasive by HT (**Fig. 4D**). Consistently, super resolution imaging also captured a portion of MT-CYB signals co-stained with DAPI as nuclear puncta in NM-treated ECs, which was remarkably increased after HT treatment (**Fig. 4E**).

**Figure 4.**
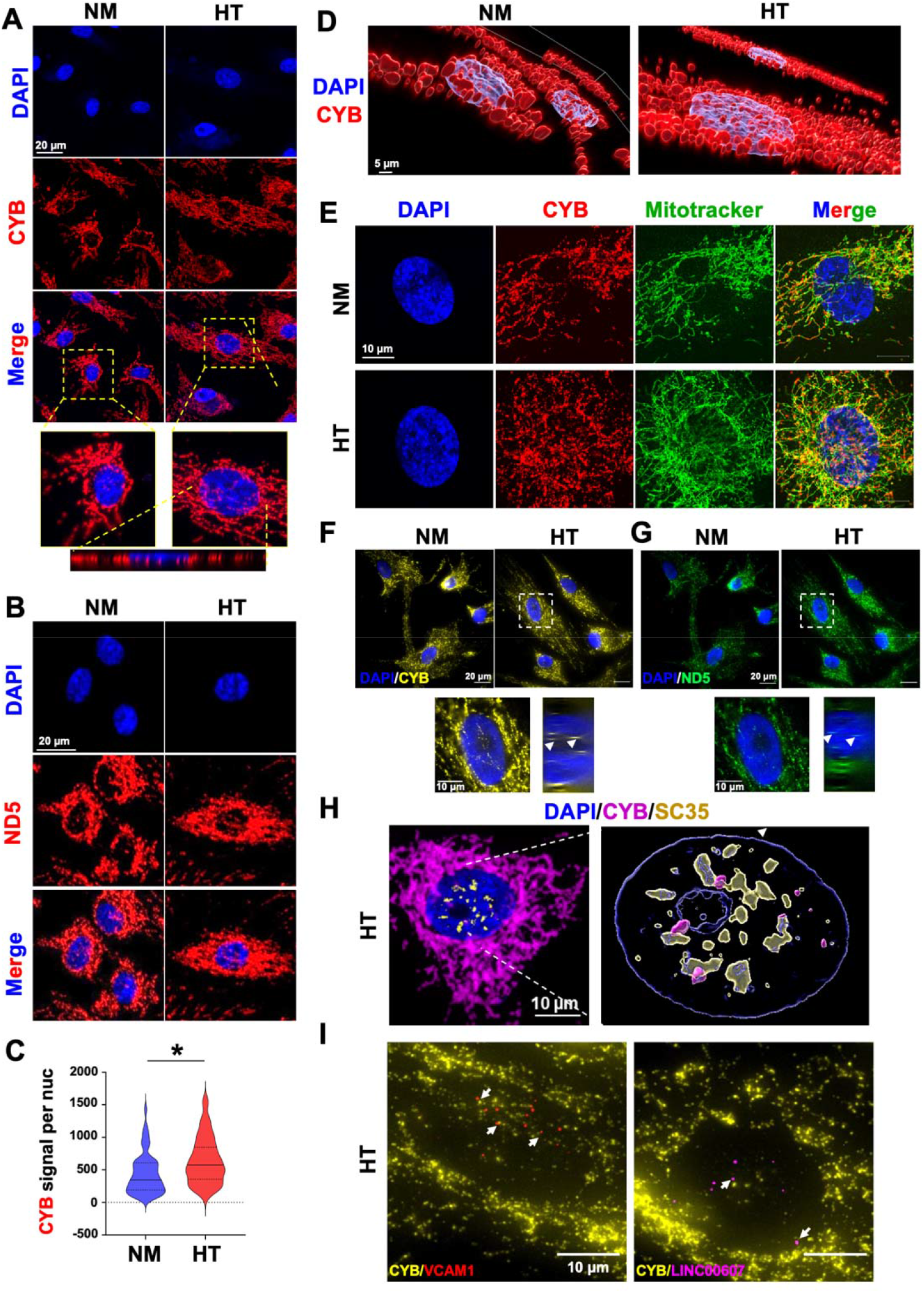
Nuclear localization of MT-CYB and MT-ND5. HUVECs were treated with NM and HT for 72 hr. (**A,B**) Confocal images of MT-CYB (A) and MT-ND5 (B) smFISH in ECs. Bottom of (A): orthogonal view across the vertical axis of HT-EC. (**C**) Violin plot showing the quantification of average MT-CYB signal per nucleus in NM and HT. Over 200 cells from each condition were quantified. * P < 0.05 by Mann-Whitney U test. (**D**) 3D reconstruction of confocal images from (A). For (A-D) representative images from three independent experiments are shown. (**E**) Super resolution microscopic images of MT-CYB smFISH and Mitotracker deep red from a single optical plane of 0.16 μm thickness. (**F,G**) Single molecule sequential FISH with probes for CYB and ND5, with DAPI staining nuclei. (**H**) Super resolution microscopic image of merged signals of smFISH for MT-CYB RNA (in magenta), immunofluorescence for SC35 protein (in yellow), and DAPI (in blue) from a single optical plane of 0.16 μm thickness (left) and the 3D-reconstruction (right). (**I**) co-sm-seqFISH for indicated pre-mRNA transcripts with MT-CYB transcript. (H) and (I) show images taken from HT-treated ECs.

As an independent validation, we designed separate pools of probes and performed sequential smFISH (*44, 45*), which yielded consistent data showing the nuclear localization of MT-CYB and MT-ND5 especially in HT-treated ECs (**Fig. 4F,G**). To test the hypothesized role of mtRNA in transcriptional activation of the nuclear genome, we performed two-fold experiments: (i) smFISH for MT-CYB and co-immunofluorescence (IF) using SC35 antibody which marks nuclear speckles and actively transcribed regions (*46, 47*) and (ii) seqFISH for MT-CYB and pre-mRNAs (using intron-targeting probes) transcriptionally induced by HT, i.e., VCAM1 and LINC00607 (using probes targeting introns). As shown in **Fig. 4H,I**, in HT-treated ECs, a fraction of MT-CYB signals overlapped those of SC35, and the signals of MT-CYB showed proximity to those of HT-induced VCAM1 and LINC00607 pre-mRNAs.

Finally, we sought for evidence that the observed phenomenon and mechanism may occur *in vivo* and is relevant to health and disease states. We performed smFISH on ECs freshly isolated from the mesenteric arteries of human donors. To correlate with the *in vitro* experiments, we selected the non-diabetic donors and those with Type 2 diabetes (T2D) according to HbA1C levels and medical record (**Fig. 5A**). Of note, the two T2D donors each had 8 and over 20 years of diabetes and were uncompliant to treatments. In agreement with data from cultured ECs, smFISH detected nuclear MT-CYB signals in ECs from all four donors. Furthermore, T2D donor-derived ECs exhibited more nuclear localized MT-CYB puncta compared to non-diabetic donors (**Fig. 5B,C**). Together, these data provide supporting evidence for the involvement of mt-caRNAs in endothelial dysfunction in the context of diabetes.

**Figure 5.**
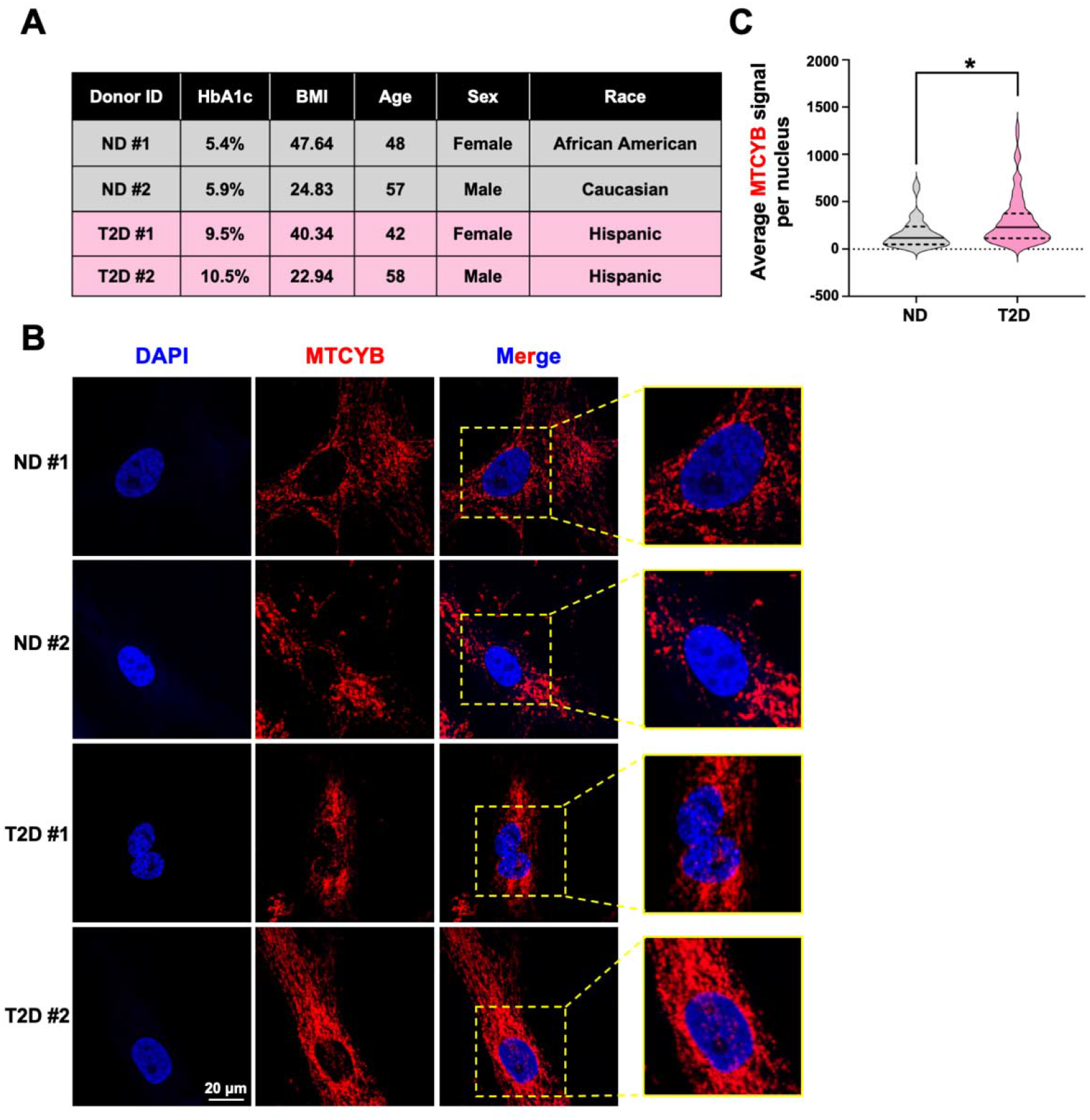
(**A-C**) Intimal cells were freshly isolated from mesenteric arteries of two non-diabetic (ND) and two Type 2 diabetic (T2D) donors and grown on glass coverslips. (**A**) Table showing donor information. (**B**) smFISH (RNAscope) was performed for MT-CYB transcripts (in red), with nuclear counter stain with DAPI. Representative images from >15 cells per donor are shown. (**C**) Violin plot showing the distribution of average MTCYB signal per nucleus from n=34 for ND and n = 81 for T2D. * P< 0.05 based on Mann-Whitney U test.

## Discussion

Mitochondrial-nuclear communication is essential for maintaining cellular homeostasis. Mitochondrial-nuclear retrograde signaling can be mediated through an array of molecules including but not limited to metabolites, peptides, and Ca^+^. These signaling molecules often activate transcription factors in the cytoplasm, which subsequently translocate into the nucleus to alter transcription. It remains an open question whether any other means of transducing retrograde signals and whether there are other types of molecules that can serve as the messenger.

To be a messenger of retrograde signaling, two criteria must be satisfied. First, this molecule must reflect certain functional status of the mitochondria. Second, this molecule must either directly or indirectly alter the epigenome and thus impact the transcription of nuclear genes. In this work, we report a widespread association of mtRNAs with nuclear chromatin. As the transcriptional output of the mitochondrial genome, mtRNA reflects the transcriptional status of the mitochondria and thus satisfies the first criterion. Furthermore, the presence of high glucose and inflammatory cytokines induced the chromatin-associated mtRNA (mt-caRNA) in human EC, confirming that the level of mt-caRNA responds to cellular stress. Mt-caRNA establishes a direct connection between mtRNA and the epigenome.

Correlated with the stress-induced increase of mt-caRNA are the transcription levels of nuclear genes including adhesion molecules ICAM1 and VCAM1, and increased monocyte-EC attachment. Depletion of a specific mt-caRNA ameliorated stress-induced transcription of nuclear genes including ICAM1 and VCAM1, and abolished the stress-induced increase in monocyte-EC attachment. Reversely, inducing this mt-caRNA in the normal condition reproduced the stress-induced transcriptional changes and the increased monocyte-EC attachment, confirming a for mt-caRNA in transcriptional regulation. These data nominate mtRNA-chromatin interaction as a means of retrograde signaling.

To ensure our discovered mt-caRNA did not originate from NUMTs, we examined the distribution of the mismatch bases between any mtRNA-aligned sequencing reads and NUMT sequences. Not only that all the several million mtRNA-aligned reads exhibited better alignment to mtRNA than any NUMTs, but most of these reads also contain 5 or more bases of mismatches to any NUMTs. These data suggest that most of mtRNA-nuDNA read pairs are not attributable to possible misalignments. Consistently, to target SncmtRNA, we used BLAST to ensure that the LNA-S GapmeR does not potentially target any homologous transcripts arising from NUMTs. Furthermore, the lack of any detectable signals for MTCYB and MTND5 in the Rho^0^ cells also excluded the potential possibility of capturing NUMT-derived transcripts in smFISH. Collectively, these controls should be sufficient to differentiate mtRNA from NUMT-derived RNA. Future studies on the mechanism of nuclear translocation of mtRNA are warranted.

Using SncmtRNA and ECs as an exemplar model to investigate the role of mt-caRNA in nuclear transcription, we observed a strong effect of Snc-KD (primarily in the nucleus) on nuclear transcriptome but not on mitochondrial gene expression and function. At basal (NM) condition, Snc-KD led to a strong suppression of anti-viral and Type 1 interferon response pathways in ECs. While similar effect was observed in HT-stressed EC, Snc-KD also suppressed multiple inflammatory response pathways. These findings support a positive regulation of mt-caRNA in innate immune and inflammatory response and are in line with the well-recognized role of mitochondria in innate immunity (*48*). In addition to the release of mtDNA and dsRNA by mitochondria into cytoplasm, mt-caRNA (exemplified by SncmtRNA) may also participate in the mitochondria-controlled intrinsic immune surveillance by transducing the defense signal to the nucleus. This mechanism may also “prime” the cells for activation of the inflammatory response under pathological stimuli, e.g. diabetes-mimicking HT or bacterial infection. Indeed, Snc-KD also attenuated LPS-induced inflammatory gene expression, including MX1, DDX58, VCAM1, and ICAM1 (**Fig. S6**). Of note, Snc-KD also had a strong effect in upregulating many cell cycle regulators, as revealed by bulk RNA-seq. However, these genes were not significant DEGs in the snRNA-seq data, suggesting that the effect of SncmtRNA on cell cycle may not be primary. It will be interesting to dissect in future studies the detailed molecular mechanisms underlying mt-caRNA-modulated nuclear transcription.

Finally, using smRNA FISH we showed that MT-CYB and MT-ND5, two other mt-caRNAs are localized in the nucleus of freshly isolated human ECs. Furthermore, their nuclear signals are increased in ECs from diabetic donors and co-localize with active transcription foci, providing *in vivo* and human disease relevance of mt-caRNAs. Although these experiments were performed for a few mtRNAs and are with limited number of human donor samples, it is likely that the chromatin attachment of mt-caRNAs is a common phenomenon present in various cell types in physiological and pathophysiological conditions, as suggested by the iMARGI data. Given the critical nuclear-mitochondria crosstalk in most cellular functions and biological processes, our study unravels a previously unidentified mechanism by which mt-caRNA forms an additional layer for nuclear transcriptional and epigenetic regulation, which may have important implications in health and disease.

## Materials and Methods

### iMARGI data analyses

iMARGI sequencing reads were processed by the iMARGI-Docker software to identify the RNA-DNA read pairs including nuRNA-nuDNA and mtRNA-nuDNA read pairs (*32*). The chromatin association level in terms of normalized CPM was obtained by dividing the number of RNA-nuDNA read pairs involving this type of RNA by the number of RNAs involved in this RNA type and by the number of total RNA-DNA read pairs in the library. The CAL of each type of RNA on any type of genomic region, e.g. promoters, was obtained by dividing the number of RNA-nuDNA read pairs involving this type of RNA and promoter DNA sequences by the total length of promoters (in kb) and by the number of RNA-nuDNA read pairs involving this type of RNA.

### Cell culture, transfection, and infection

HUVECs were cultured in complete M199 medium supplemented with 20% FBS (Sigma, F1141) and 1x antibiotics (Penicillin-Streptomycin, Gibco, 15140122). For osmolarity control (NM) samples, 25mM D-mannitol (Fisher Scientific, M120-500) was added. For HT-treated samples, 25mM D-glucose + 5ng/ml TNFα (ThermoFisher Scientific, PHC3015) were added. HEK-293 and Rho^0^ cells were cultured with DMEM (Corning 10-013-CV) supplemented with 10% FBS. For imaging experiments, ECs were cultured on coverslips (Bioptechs) pre-coated with 10mg/ml fibronectin (Sigma F2006) and 0.01% poly-L-lysine (Millipore, A-005-C).

The SncmtRNA full-length cDNA sequence was synthesized by Genscript Inc. and subcloned into pCMV-MCS packaging vector using ClaI and BglII sites. The control AAV vector, pAAV-CAG-tdTomato (Plasmid #59462) was a gift from Edward Boyden and obtained from Addgene (Addgene plasmid # 59462). HEK293A cells were co-transfected with RepCap, pHelper, and the SncmtRNA/tdTomato plasmids with PEI at 80% confluency. The cells and media were harvested after 96 hrs and centrifuged at 1000 rpm for 10 min at 4°C. The pellets were resuspended in PBS containing 0.0001% pluromic F68 and 200 mM NaCl and freeze-thawed in liquid nitrogen. The cell debris were pelleted at 3220g for 15 min at 4°C and the supernatant containing the cleared lysate and virus was filtered through a 0.45 μm PES membrane before resuspending in PEG solution.

The IRF3/IRF7 luciferase reporter was constructed by subcloning 3x IRF3/7 binding element (GTCAGGAGAAGGAAACCTTC) into the Sal I and HindIII sites in the pGL3-basic (Promega E1751) backbone vector. HEK-293 cells were co-transfected with Renilla and IRF3/7 using Lipofectamine 2000 (Thermo Fisher scientific) following manufacturer’s protocol. After 48 hrs, cells were lysed using Reporter Lysis Buffer (Promega) and the luciferase activity was measured using the Luciferase Assay System (Promega, Cat # E4030) following manufacturer guidelines.

### FISH for SncmtRNA

FISH for SncmtRNA was performed using a FAM-labelled LNA-S probe (sequence shown in **Table S1**) following Stellaris RNA FISH protocol for adherent cells available online at www.biosearchtech.com/stellarisprotocolsth with slight modifications. A FAM-labelled LNA probe with scrambled sequence was used a control. Briefly, cells were grown and fixed on coverslips using 4% PFA. Cells were permeabilized in 70% ethanol for 1 hr followed by hybridization with the probe (200 nM final concentration) in the Stellaris hybridization buffer overnight at 37°C. Subsequently, cells were washed with Stellaris wash buffers A and B, stained with DAPI, and mounted for imaging. Images were taken on Zeiss LSM 880 in Airyscan mode.

### RNA extraction and quantitative PCR

Total RNA was isolated from cells using TRIzol according to manufacturer’s protocol. TURBO DNase (ThermoFisher Scientific, AM2238) was used to eliminate genomic DNA before reverse transcription with SuperScript^™^ IV Reverse Transcriptase (ThermoFisher Scientific, 18090010) using random hexamer primers. qPCR was performed with Bio-Rad SYBR Green Supermix on Bio-Rad CFX Connect Real Time system. For detecting MT-RNR2-derived transcripts, the extension time per cycle was increased from the standard 30 sec to 45 sec. All primer sequences used for qPCR amplification are listed in **Table 1**.

### Subcellular Fractionation

To isolate RNA from the nuclear and cytoplasmic fractions, HUVECs were collected in 1X PBS using a cell scraper. The cells were pelleted at 300g for 10 min, resuspended in nuclear isolation buffer (1.28 M sucrose, 40 mM Tris-HCl pH 7.5, 20 mM MgCl_2_, 4% Triton X-100, 1X protease inhibitor, 1X PMSF, and recombinant RNase inhibitor), and incubated on ice for 20 mins while vortexing every 5 min. After 20 min, the cell lysates were centrifuged at 2500g for 15 mins in a swing bucket centrifuge. The supernatant containing the cytoplasmic fraction and the pellet containing the nuclear fraction were used to isolate RNA using TRIzol LS and TRIzol respectively. RNA isolation from nucleoplasmic and chromatin fraction was performed as described previously (*34*).

To isolate RNA from the mitochondrial fraction, we followed a protocol described by Dhir et al. (*21*) using MACS mitochondrial isolation kit (Miltenyi Biotec, #130-094-532). Briefly, cell pellets were resuspended in 500 μl lysis buffer provided with the kit, homogenized using a 25G needle, made up to 10 ml with 1X separation buffer, and incubated with 50 μl TOM-22 beads for 1hr with rotation at 4°C. Following the incubation, the bound fractions were eluted into 1ml separation buffer using the MACS LS columns and centrifuged at 14000g for 15 mins, followed by RNA extraction using TRIzol.

### Bulk RNA sequencing and analysis

Total RNA libraries were prepared using the KAPA mRNA HyperPrep Kit (Roche Diagnostics) following the manufacturer’s manual. For analysis, raw sequencing reads were processed using Trim Galore and the trimmed reads were aligned to the hg38 reference genome using HISAT2 (*49*). HTseq (*50*) was used to generate counts and DESeq2 (*51*) was then used to perform differential gene expression analysis with default parameters (genes with P values < 0.05 and Log_2_ fold-change > |0.5| were considered significant differently expressed genes).

### Monocyte adhesion assay

Leukapheresis products and discards were obtained from consented research participants (healthy donors) under protocols approved by the City of Hope Internal Review Board (IRB #09025). Peripheral blood mononuclear cells (PBMC) were isolated by density gradient centrifugation over Ficoll-Paque (GE Healthcare) followed by multiple washes in PBS/EDTA (Miltenyi Biotec). Cells were rested overnight at room temperature on a rotator, and subsequently washed and resuspended in complete X-VIVO. Up to 5×10^9^ PBMC were incubated with anti-CD14 microbeads (Miltenyi Biotec) for 30 minutes at room temperature and magnetically enriched using the CliniMACS system (Miltenyi Biotec) according to the manufacturer’s protocol. The monocytes were labeled with CellTracker™ Green CMFDA Dye (Thermo Fisher Scientific) and incubated with monolayer HUVECs (4□×□10^3^ cells per cm^2^) for 15–30 min in a cell culture incubator. The nonattached monocytes were then washed off with complete EC growth medium. The attached monocyte numbers were quantified on Cytation^™^1 Cell Imaging Multi-Mode Reader (BioTek) using green fluorescent channel. Average numbers per condition were calculated from two randomly selected fields of technical duplicates.

### Fluorescence ubiquitination cell cycle indicator (FUCCI) analysis

Cell cycle analysis was performed using the Premo FUCCI Cell Cycle Sensor kit (Thermo Fisher Scientific; Cat # P36237) following the manufacturer’s protocol. Briefly, Premo geminin-GFP and Premo Cdt1-RFP were added to the EC culture medium at 80 particles per cell. Cdt1 levels are highest during G1, while Geminin levels are highest during the S, G2, and M phases. Thus, tracking the fluorescence will enable monitoring cell cycle progression overtime. Twenty-four hr later, cells were imaged using Cytation™1 Cell Imaging Multi-Mode Reader (BioTek). Green fluorescent channel was used for GFP, and red fluorescent channel for RFP. The number of cells with red fluorescence (indicative of G1 phase), yellow fluorescence (indicative of G1/S transition), and green fluorescence (indicative of G2/M) were counted from at least 4 fields per condition and five independent experiments.

### scRNA- and snRNA-sequencing and analysis

HUVECs treated with HT and transfected with LNA in biological replicates were processed following the Drop-seq protocol from 10x Genomics. For scRNA-seq, ECs were washed, trypsinized, and suspended into a single-cell solution for the library preparation using 10x Genomics Chromium 3.1’ expression kit as previously described (*34*). For snRNA-seq, cells were first washed with 1X PBS+ 0.04% BSA, pelleted at 300g for 5 min, and resuspended in 100 μl of freshly prepared lysis buffer (10 mM Tris-HCl pH 7.4, 10 mM NaCl, 3 mM MgCl2, 0.1% Tween-20, 0.1% Nonidet P40 Substitute, 0.01% Digitonin, 1% BSA). The cells were then lysed on ice for 5 min, checked for nuclear integrity under microscope, and washed 3 times in wash buffer (10 mM Tris-HCl pH 7.4, 10 mM NaCl, 3 mM MgCl_2_, 0.1% Tween-20, 1% BSA). The resulting nuclei underwent the Drop-seq protocol with 10x Genomics Chromium 3.1’ expression kit.

scRNA-seq and snRNA-seq data were processed using the standardized pipeline provided by 10x Genomics. The reads were aligned to both intronic and exonic regions of the human hg38 reference genome by utilizing include-introns command during alignment with Cell Ranger. Seurat R package (v3) was used following well-established filtering steps to remove genes expressed in < 3 cells and cells expressing < 200 genes. For the differential analysis between HT-Scr and HT-LNA conditions, rare cells/nuclei with very high numbers of genes (> 8,500 and 7,000 genes for scRNA-seq and snRNA-seq respectively; potential multiplets) and cells/nuclei with high mitochondrial read percentages (> 15% for scRNA-seq and >25% for snRNA-seq) were removed. HT_Scr and HT_LNA samples were first separately normalized with “sctransform” and then integrated into one dataset based on Seurat-selected anchor genes. This integrated dataset was used for clustering analysis. Clusters of nuclei were identified by Seurat based on shared nearest neighbor (SNN) using the first 10 principal components as input (resolution = 0.5 Differential expression analysis was performed using the Wilcoxon test with default parameters in Seurat. The threshold for the avg_logFC was set to be 0.1.

### Nascent RNA pull-down

Nascent RNA pull-down assay was performed as previously reported (*34*). Briefly, ECs were synchronized by serum starvation with M199 + 2% FBS for 6 hrs. Nascent/newly synthesized RNA were labelled with 5-Ethynyl Uridine (EU) at 200 μM final concentration and incubated for 24 hr. Total RNA was then isolated from the cells and 3 μg of total RNA was used to capture nascent RNA using the Click-iT Nascent RNA Capture Kit (Thermo Fisher Scientific, C10365) following manufacturer’s protocol.

### Single-molecule Fluorescence *in situ* hybridization (smFISH), co-smFISH-Immunofluorescence (IF), and confocal and super-resolution microscopy

smFISH shown in Fig. 4A-E,H and Fig. S5 were performed using RNAscope™ Multiplex Fluorescent V2 Assay (ACDBio, 323100) following manufacturer’s protocol. Briefly, cells were fixed with 4% paraformaldehyde (PFA) for 30 min at room temperature, ethanol-dehydrated, pre-treated with hydrogen peroxide (H_2_O_2_) for 10 min at room temperature, and permeabilized with Protease III (1:10 dilution) for 30 min at room temperature prior to probe hybridization following recommended protocol. RNAscope^®^ Probe - Hs-MT-CYB (Cat # 582771), RNAscope^®^ Probe - Hs-MT-ND5-C2 (Cat # 539451-C2) were used to detect human mitochondrial MT-CYB and MT-ND5 transcripts respectively. Following hybridization, the RNAscope assay was developed following the manufacturer’s protocol. Opal dyes 570 or 690 were used to detect the mitochondrial transcripts. To image mitochondria and mitochondrial transcripts simultaneously, cells were incubated with MitoTracker™ Deep Red FM (Thermo Fisher Scientific, M22426) at 100 nM final concentration in serum-free media for 30 min prior to fixing.

smFISH combined with IF was performed using the RNA-Protein Co-Detection Ancillary Kit (ACDBio, Cat # 323180) along with the RNAscope™ Multiplex Fluorescent V2 Assay (ACDBio, Cat # 323100) following the manufacturer protocol. Briefly, cells were fixed as in smFISH and blocked with co-detection antibody diluent for 30 min at room temperature prior to incubating with anti-human SC35 (Abcam, Cat # ab11826) at 1:50 dilution overnight at 4°C. Cells were washed twice with 1X PBST (1X PBS Ca^2+^ and Mg^2+^ free + 0.1% Tween-20) and fixed again with 4% PFA before proceeding with permeabilization and RNAscope assay as abovementioned. After completing the final step of the RNAscope assay, the cells were incubated with secondary antibody anti-mouse Alexa 488 (Thermo Fisher Scientific, Cat # A-11008) at 1:200 dilution for 30 min at room temperature, washed twice with 1X PBST and mounted for imaging. DAPI was used to stain the DNA in the nuclei.

Confocal microscopy was performed on Zeiss LSM 700 microscope using 20X or 63X water immersion lens. 3D reconstruction of confocal images was performed using Imaris software with “Surfaces Technology”. Super-resolution images were captured on Zeiss LSM 880 microscope using 63X oil immersion lens in the Airyscan mode.

Image analysis for smFISH was performed using FIJI (ImageJ software) in a blinded fashion. A mask was drawn around the nuclear region (as defined by DAPI channel), and the average gray value of MTCYB signals within this mask was used to compute MTCYB signals per nucleus.

smFISH shown in **Fig. 4F,I** were performed following a previously published protocol (ref) with modifications. The probes were designed using ProbeDealer (*52*) and synthesized by IDT. ProbeDealer design is based on melting temperature, GC content, exclusion of repetitive sequences and the specificity was further confirmed by BLAST. Each probe consists of a 30-nt target region that specifically binds to transcript of interest and a 20-nt readout region that allows the binding of dye-labeled secondary oligos as previously validated (*45, 53*). Specifically, 48 oligos for VCAM and LINC00607 and 32 oligos for MT-CYB and 45 oligos for MT-ND5 were used.

Cells were fixed with 4% PFA (EMS, 15710), permeabilized with 0.5% Triton X-100 (Sigma, T8787), and incubated in the pre-hybridization buffer with 50% formamide (Sigma, F7503) in 2x saline-sodium citrate (SSC, Invitrogen 15557-044) for 5 min at RT and then in the hybridization buffer containing 50% formamide, 0.1% yeast tRNA (Life Technologies, 15401011), 10% dextran sulfate (Millipore, S4030) and 1% murine RNase inhibitor (New England Biolabs, M0314L) in 2× SSC with 6 nM primary probes at a at 37°C for 24 hr. Samples were then washed with 2×SSC containing 0.1% Tween 20 at 60□°C, followed by 2x SSCT wash at RT. The sample then underwent multiple rounds of sequential imaging, using Translura’s automated sequential smFISH microscope, and corresponding secondary hybridization, washing and imaging buffers. Final concentration of the secondary oligos were 3.75nM. Z-stack images were taken with 750-nm or 647-nm laser, with a step size of 600nm and total Z-stacks range of 10μm.

### Donor-derived ECs

Human mesenteric artery tissue studies were conducted on deidentified specimens obtained from the Southern California Islet Cell Resource Center at City of Hope. The research consents for the use of postmortem human tissues were obtained from the donors’ next of kin and ethical approval for this study was granted by the Institutional Review Board of City of Hope (IRB #01046). Type 2 Diabetes (T2D) was identified based on diagnosis in the donors’ medical records and the percentage of glycated hemoglobin A1c (HbA1c) of 6.5% or higher. Donor-derived ECs were isolated following our published protocol (*54*). Briefly, the mesenteric arteries were cut open lengthwise to expose the lumen. The intimal layer containing ECs was scraped mechanically using a scalpel in pre-warmed digestion buffer containing Collagenase D. The collected intimal cells were digested for 5 min at 37°C, washed with complete M199 medium, and then plated onto tissue culture dishes coated with attachment factor or collagen. Upon cell attachment and growth, the cells were transferred onto a cover glass for smFISH experiments.

### Statistical analysis

For all experiments, at least 3 independent experiments were performed. Statistical analysis was performed using Student t-tests for two group comparisons or ANOVA with Bonferroni post-hoc test for multiple group comparisons. For all the high-throughput sequencing data, experiments were performed in 2-3 biological replicates. P < 0.05 was considered statistically significant unless otherwise indicated in the figure legends.

## Acknowledgments

The authors thank Dr. John Burnett at City of Hope for providing Rho^0^ cells, Drs. Ismail Al-Abdullah and Meirigeng Qi of the islet transplantation team at City of Hope for isolation of human arteries, Drs. John Y-J. Shyy, John Rossi, Yilun Liu, and John Burnett for helpful discussion and valuable inputs, and Dr. Siyuan Wang at Yale University for technical consultation on smFISH.

## Funding

National Institutes of Health grant R01 HL145170 (ZBC)

National Institutes of Health grants R01GM138852, R01HD107206, and DP1DK126138 (SZ)

National Institutes of Health grant R01 HL106089 to RN and ZBC

Ella Fitzgerald Foundation (ZBC)

Research National Institutes of Health P30CA033572 supported work performed in the Integrative Genomics, Light Microscopy and Digital Imaging Cores

## Author contributions

Conceptualization: KS, SZ, ZBC

Methodology: KS, ZQ, NKM, XL, YL, MS, SJ, JS

Investigation: KS, ZQ, RC, DQ, NKM, XL, YL

Supervision: BA, RN, SZ, ZBC

Vector construction: RD, JL

Writing-original draft: KS, ZQ, SZ, ZBC

Writing-review & editing: KS, ZQ, SZ, ZBC, SJP, PW, RN, DQ, XL, YL, NKM

## Data and materials availability

All data needed to evaluate the conclusions in the paper are present in the paper and/or the Supplementary Materials and all high-throughput sequencing data used in this study have been deposited at GEO with accession No. GSE211971.

## References

1. D. C. Wallace, A mitochondrial paradigm of metabolic and degenerative diseases, aging, and cancer: a dawn for evolutionary medicine. Annu Rev Genet 39, 359–407 (2005).

2. Tim. R. Mercer, S. Neph, Marcel E. Dinger, J. Crawford, Martin A. Smith, A.-Marie. J. Shearwood, E. Haugen, Cameron P. Bracken, O. Rackham, John A. Stamatoyannopoulos, A. Filipovska, John S. Mattick, The Human Mitochondrial Transcriptome. Cell 146, 645–658 (2011).

3. O. Rackham, A.-M. J. Shearwood, T. R. Mercer, S. M. K. Davies, J. S. Mattick, A. Filipovska, Long noncoding RNAs are generated from the mitochondrial genome and regulated by nuclear-encoded proteins. RNA 17, 2085–2093 (2011).

4. S. Ro, H. Y. Ma, C. Park, N. Ortogero, R. Song, G. W. Hennig, H. Zheng, Y. M. Lin, L. Moro, J. T. Hsieh, W. Yan, The mitochondrial genome encodes abundant small noncoding RNAs. Cell Res 23, 759–774 (2013).

5. P. M. Quiros, A. Mottis, J. Auwerx, Mitonuclear communication in homeostasis and stress. Nat Rev Mol Cell Biol 17, 213–226 (2016).

6. N. Gleyzer, K. Vercauteren, R. C. Scarpulla, Control of mitochondrial transcription specificity factors (TFB1M and TFB2M) by nuclear respiratory factors (NRF-1 and NRF-2) and PGC-1 family coactivators. Mol Cell Biol 25, 1354–1366 (2005).

7. J. V. Virbasius, R. C. Scarpulla, Activation of the human mitochondrial transcription factor A gene by nuclear respiratory factors: a potential regulatory link between nuclear and mitochondrial gene expression in organelle biogenesis. Proc Natl Acad Sci U S A 91, 1309–1313 (1994).

8. D. Ojala, J. Montoya, G. Attardi, tRNA punctuation model of RNA processing in human mitochondria. Nature 290, 470–474 (1981).

9. M. Falkenberg, M. Gaspari, A. Rantanen, A. Trifunovic, N. G. Larsson, C. M. Gustafsson, Mitochondrial transcription factors B1 and B2 activate transcription of human mtDNA. Nat Genet 31, 289–294 (2002).

10. I. Kuhl, M. Miranda, V. Posse, D. Milenkovic, A. Mourier, S. J. Siira, N. A. Bonekamp, U. Neumann, A. Filipovska, P. L. Polosa, C. M. Gustafsson, N. G. Larsson, POLRMT regulates the switch between replication primer formation and gene expression of mammalian mtDNA. Sci Adv 2, e1600963 (2016).

11. R. Vendramin, Y. Verheyden, H. Ishikawa, L. Goedert, E. Nicolas, K. Saraf, A. Armaos, R. Delli Ponti, K. Izumikawa, P. Mestdagh, D. L. J. Lafontaine, G. G. Tartaglia, N. Takahashi, J. C. Marine, E. Leucci, SAMMSON fosters cancer cell fitness by concertedly enhancing mitochondrial and cytosolic translation. Nat Struct Mol Biol 25, 1035–1046 (2018).

12. R. S. Puranam, G. Attardi, The RNase P associated with HeLa cell mitochondria contains an essential RNA component identical in sequence to that of the nuclear RNase P. Mol Cell Biol 21, 548–561 (2001).

13. X. Zhang, X. Zuo, B. Yang, Z. Li, Y. Xue, Y. Zhou, J. Huang, X. Zhao, J. Zhou, Y. Yan, H. Zhang, P. Guo, H. Sun, L. Guo, Y. Zhang, X.-D. Fu, MicroRNA Directly Enhances Mitochondrial Translation during Muscle Differentiation. Cell 158, 607–619 (2014).

14. D. D. Stefani, R. Rizzuto, T. Pozzan, Enjoy the Trip: Calcium in Mitochondria Back and Forth. Annual Review of Biochemistry 85, 161–192 (2016).

15. Z. Liu, R. A. Butow, Mitochondrial Retrograde Signaling. Annual Review of Genetics 40, 159–185 (2006).

16. M. G. Vizioli, T. Liu, K. N. Miller, N. A. Robertson, K. Gilroy, A. B. Lagnado, A. Perez-Garcia, C. Kiourtis, N. Dasgupta, X. Lei, P. J. Kruger, C. Nixon, W. Clark, D. Jurk, T. G. Bird, J. F. Passos, S. L. Berger, Z. Dou, P. D. Adams, Mitochondria-to-nucleus retrograde signaling drives formation of cytoplasmic chromatin and inflammation in senescence. Genes & Development 34, 428–445 (2020).

17. J. C. Reynolds, R. W. Lai, J. S. T. Woodhead, J. H. Joly, C. J. Mitchell, D. Cameron-Smith, R. Lu, P. Cohen, N. A. Graham, B. A. Benayoun, T. L. Merry, C. Lee, MOTS-c is an exercise-induced mitochondrial-encoded regulator of age-dependent physical decline and muscle homeostasis. Nature Communications 12, 470 (2021).

18. L. S. Huang, Z. Hong, W. Wu, S. Xiong, M. Zhong, X. Gao, J. Rehman, A. B. Malik, mtDNA Activates cGAS Signaling and Suppresses the YAP-Mediated Endothelial Cell Proliferation Program to Promote Inflammatory Injury. Immunity 52, 475–486.e475 (2020).

19. H. Maekawa, T. Inoue, H. Ouchi, T. M. Jao, R. Inoue, H. Nishi, R. Fujii, F. Ishidate, T. Tanaka, Y. Tanaka, N. Hirokawa, M. Nangaku, R. Inagi, Mitochondrial Damage Causes Inflammation via cGAS-STING Signaling in Acute Kidney Injury. Cell Rep 29, 1261–1273 e1266 (2019).

20. A. P. West, W. Khoury-Hanold, M. Staron, M. C. Tal, C. M. Pineda, S. M. Lang, M. Bestwick, B. A. Duguay, N. Raimundo, D. A. MacDuff, S. M. Kaech, J. R. Smiley, R. E. Means, A. Iwasaki, G. S. Shadel, Mitochondrial DNA stress primes the antiviral innate immune response. Nature 520, 553–557 (2015).

21. A. Dhir, S. Dhir, L. S. Borowski, L. Jimenez, M. Teitell, A. Rötig, Y. J. Crow, G. I. Rice, D. Duffy, C. Tamby, T. Nojima, A. Munnich, M. Schiff, C. R. de Almeida, J. Rehwinkel, A. Dziembowski, R. J. Szczesny, N. J. Proudfoot, Mitochondrial double-stranded RNA triggers antiviral signalling in humans. Nature 560, 238–242 (2018).

22. M. Tigano, D. C. Vargas, S. Tremblay-Belzile, Y. Fu, A. Sfeir, Nuclear sensing of breaks in mitochondrial DNA enhances immune surveillance. Nature 591, 477–481 (2021).

23. J. Villegas, V. Burzio, C. Villota, E. Landerer, R. Martinez, M. Santander, R. Martinez, R. Pinto, M. I. Vera, E. Boccardo, L. L. Villa, L. O. Burzio, Expression of a novel noncoding mitochondrial RNA in human proliferating cells. Nucleic Acids Research 35, 7336–7347 (2007).

24. V. A. Burzio, C. Villota, J. Villegas, E. Landerer, E. Boccardo, L. L. Villa, R. Martínez, C. Lopez, F. Gaete, V. Toro, X. Rodriguez, L. O. Burzio, Expression of a family of noncoding mitochondrial RNAs distinguishes normal from cancer cells. Proceedings of the National Academy of Sciences 106, 9430–9434 (2009).

25. E. Landerer, J. Villegas, V. A. Burzio, L. Oliveira, C. Villota, C. Lopez, F. Restovic, R. Martinez, O. Castillo, L. O. Burzio, Nuclear localization of the mitochondrial ncRNAs in normal and cancer cells. Cellular Oncology 34, 297–305 (2011).

26. A. Blumental-Perry, R. Jobava, I. Bederman, A. J. Degar, H. Kenche, B. J. Guan, K. Pandit, N. A. Perry, N. D. Molyneaux, J. Wu, E. Prendergas, Z. W. Ye, J. Zhang, C. E. Nelson, F. Ahangari, D. Krokowski, S. H. Guttentag, P. A. Linden, D. M. Townsend, A. Miron, M. J. Kang, N. Kaminski, Y. Perry, M. Hatzoglou, Retrograde signaling by a mtDNA-encoded non-coding RNA preserves mitochondrial bioenergetics. Communications Biology 3, 626 (2020).

27. X. Li, X. D. Fu, Chromatin-associated RNAs as facilitators of functional genomic interactions. Nat Rev Genet 20, 503–519 (2019).

28. T. C. Nguyen, K. Zaleta-Rivera, X. Huang, X. Dai, S. Zhong, RNA, Action through Interactions. Trends Genet 34, 867–882 (2018).

29. X. W. Riccardo Calandrelli, John L Charles Richard, Zhifei Luo, Tri C. Nguyen, Chien-Ju Chen, Zhijie Qi, Shuanghong Xue, Weizhong Chen, Zhangming Yan, Weixin Wu, Kathia Zaleta-Rivera, Rong Hu, Miao Yu, Yuchuan Wang, Wenbo Li, Jian Ma, Bing Ren, Sheng Zhong, Three-dimensional organization of chromatin associated RNAs and their role in chromatin architecture in human cells. BioRxiv 2021.06.10.447969, (2021).

30. Z. Yan, N. Huang, W. Wu, W. Chen, Y. Jiang, J. Chen, X. Huang, X. Wen, J. Xu, Q. Jin, K. Zhang, Z. Chen, S. Chien, S. Zhong, Genome-wide colocalization of RNA-DNA interactions and fusion RNA pairs. Proceedings of the National Academy of Sciences 116, 3328–3337 (2019).

31. B. Sridhar, M. Rivas-Astroza, T. C. Nguyen, W. Chen, Z. Yan, X. Cao, L. Hebert, S. Zhong, Systematic Mapping of RNA-Chromatin Interactions In Vivo. Curr Biol 27, 602–609 (2017).

32. W. Wu, Z. Yan, T. C. Nguyen, Z. Bouman Chen, S. Chien, S. Zhong, Mapping RNA–chromatin interactions by sequencing with iMARGI. Nature Protocols 14, 3243–3272 (2019).

33. J. V. Lopez, N. Yuhki, R. Masuda, W. Modi, S. J. O’Brien, Numt, a recent transfer and tandem amplification of mitochondrial DNA to the nuclear genome of the domestic cat. J Mol Evol 39, 174–190 (1994).

34. R. Calandrelli, L. Xu, Y. Luo, W. Wu, X. Fan, T. Nguyen, C.-J. Chen, K. Sriram, X. Tang, A. B. Burns, R. Natarajan, Z. B. Chen, S. Zhong, Stress-induced RNA-chromatin interactions promote endothelial dysfunction. Nature Communications 11, 5211 (2020).

35. N. S. Chandel, P. T. Schumacker, Cells depleted of mitochondrial DNA (rho0) yield insight into physiological mechanisms. FEBS Lett 454, 173–176 (1999).

36. S. M. Shenouda, M. E. Widlansky, K. Chen, G. Xu, M. Holbrook, C. E. Tabit, N. M. Hamburg, A. A. Frame, T. L. Caiano, M. A. Kluge, M.-A. Duess, A. Levit, B. Kim, M.-L. Hartman, L. Joseph, O. S. Shirihai, J. A. Vita, Altered Mitochondrial Dynamics Contributes to Endothelial Dysfunction in Diabetes Mellitus. Circulation 124, 444–453 (2011).

37. K. Trudeau, A. J. Molina, W. Guo, S. Roy, High glucose disrupts mitochondrial morphology in retinal endothelial cells: implications for diabetic retinopathy. Am J Pathol 177, 447–455 (2010).

38. X. H. Liang, H. Sun, J. G. Nichols, S. T. Crooke, RNase H1-Dependent Antisense Oligonucleotides Are Robustly Active in Directing RNA Cleavage in Both the Cytoplasm and the Nucleus. Mol Ther 25, 2075–2092 (2017).

39. A. Sakaue-Sawano, H. Kurokawa, T. Morimura, A. Hanyu, H. Hama, H. Osawa, S. Kashiwagi, K. Fukami, T. Miyata, H. Miyoshi, T. Imamura, M. Ogawa, H. Masai, A. Miyawaki, Visualizing spatiotemporal dynamics of multicellular cell-cycle progression. Cell 132, 487–498 (2008).

40. W. Wu, H. Xiao, A. Laguna-Fernandez, G. Villarreal, Jr., K. C. Wang, G. G. Geary, Y. Zhang, W. C. Wang, H. D. Huang, J. Zhou, Y. S. Li, S. Chien, G. Garcia-Cardena, J. Y. Shyy, Flow-Dependent Regulation of Kruppel-Like Factor 2 Is Mediated by MicroRNA-92a. Circulation 124, 633–641 (2011).

41. Y. Fang, P. F. Davies, Site-specific microRNA-92a regulation of Kruppel-like factors 4 and 2 in atherosusceptible endothelium. Arterioscler Thromb Vasc Biol 32, 979–987 (2012).

42. D. Wang, P. W. L. Tai, G. Gao, Adeno-associated virus vector as a platform for gene therapy delivery. Nat Rev Drug Discov 18, 358–378 (2019).

43. F. Wang, J. Flanagan, N. Su, L. C. Wang, S. Bui, A. Nielson, X. Wu, H. T. Vo, X. J. Ma, Y. Luo, RNAscope: a novel in situ RNA analysis platform for formalin-fixed, paraffin-embedded tissues. J Mol Diagn 14, 22–29 (2012).

44. A. Raj, P. van den Bogaard, S. A. Rifkin, A. van Oudenaarden, S. Tyagi, Imaging individual mRNA molecules using multiple singly labeled probes. Nat Methods 5, 877–879 (2008).

45. M. Liu, Y. Lu, B. Yang, Y. Chen, J. S. D. Radda, M. Hu, S. G. Katz, S. Wang, Multiplexed imaging of nucleome architectures in single cells of mammalian tissue. Nat Commun 11, 2907 (2020).

46. S. Lin, G. Coutinho-Mansfield, D. Wang, S. Pandit, X. D. Fu, The splicing factor SC35 has an active role in transcriptional elongation. Nat Struct Mol Biol 15, 819–826 (2008).

47. J. Kim, N. C. Venkata, G. A. Hernandez Gonzalez, N. Khanna, A. S. Belmont, Gene expression amplification by nuclear speckle association. J Cell Biol 219, (2020).

48. A. P. West, G. S. Shadel, S. Ghosh, Mitochondria in innate immune responses. Nature Reviews Immunology 11, 389–402 (2011).

49. M. Pertea, D. Kim, G. M. Pertea, J. T. Leek, S. L. Salzberg, Transcript-level expression analysis of RNA-seq experiments with HISAT, StringTie and Ballgown. Nat Protoc 11, 1650–1667 (2016).

50. S. Anders, P. T. Pyl, W. Huber, HTSeq--a Python framework to work with high-throughput sequencing data. Bioinformatics 31, 166–169 (2015).

51. M. I. Love, W. Huber, S. Anders, Moderated estimation of fold change and dispersion for RNA-seq data with DESeq2. Genome Biol 15, 550 (2014).

52. M. Hu, B. Yang, Y. Cheng, J. S. D. Radda, Y. Chen, M. Liu, S. Wang, ProbeDealer is a convenient tool for designing probes for highly multiplexed fluorescence in situ hybridization. Sci Rep 10, 22031 (2020).

53. K. H. Chen, A. N. Boettiger, J. R. Moffitt, S. Wang, X. Zhuang, RNA imaging. Spatially resolved, highly multiplexed RNA profiling in single cells. Science 348, aaa6090 (2015).

54. N. K. Malhi, Y. Luo, X. Tang, K. Sriram, R. Calandrelli, S. Zhong, Z. B. Chen, Isolation and Profiling of Human Primary Mesenteric Arterial Endothelial Cells at the Transcriptome Level. J Vis Exp, (2022).

