## Supplemental material for "Regulation of Nuclear Transcription by Mitochondrial RNA"

Sriram *et al.*

* Co-corresponding authors

Sheng Zhong, PhD

Zhen B. Chen, PhD

**This PDF file includes:**

Figs. S1 to S6
Table S1

Fig. S1.

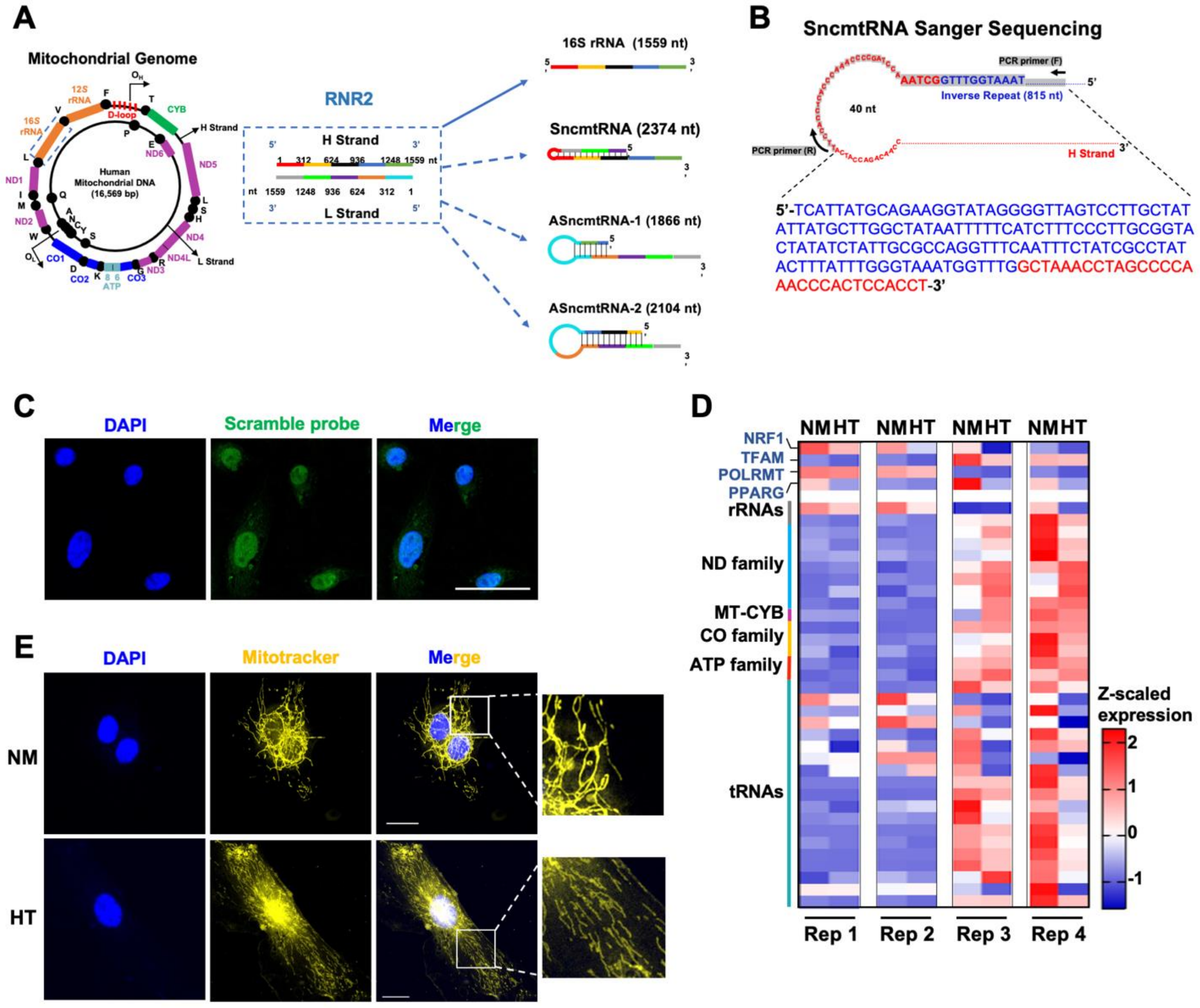


***MT-RNR2*-derived SncmtRNA and HT-regulated mitochondrial morphology and transcription.** **(A)** Schematic showing *MT-RNR2*-derived transcripts including 16S rRNA, SncmtRNA, ASncmtRNA-1, and ASncmtRNA-2. **(B)** Sanger sequencing of RT-PCR product from ECs using the indicated primers of SncmtRNA confirmed the putative chimeric junction. **(C)** Representative FISH images performed with a FAM-labeled Scramble probe in HUVECs. Scale bar = 50 µm. **(D)** Expression of nuclear encoded genes regulating mitochondrial biogenesis (*NRF1, TFAM, POLRMT,* and *PPARG*) and those transcribed from mitochondrial genome, identified by bulk RNA-seq of ECs subjected to NM or HT for 72 hrs in four biological replicates. **(E)** Mitotracker deep red staining (yellow pseudocolor) of ECs treated with NM or HT, with DAPI staining the nucleus. Representative images from three independent experiments are shown. Scale bar = 20 µm.

**Fig. S2.**

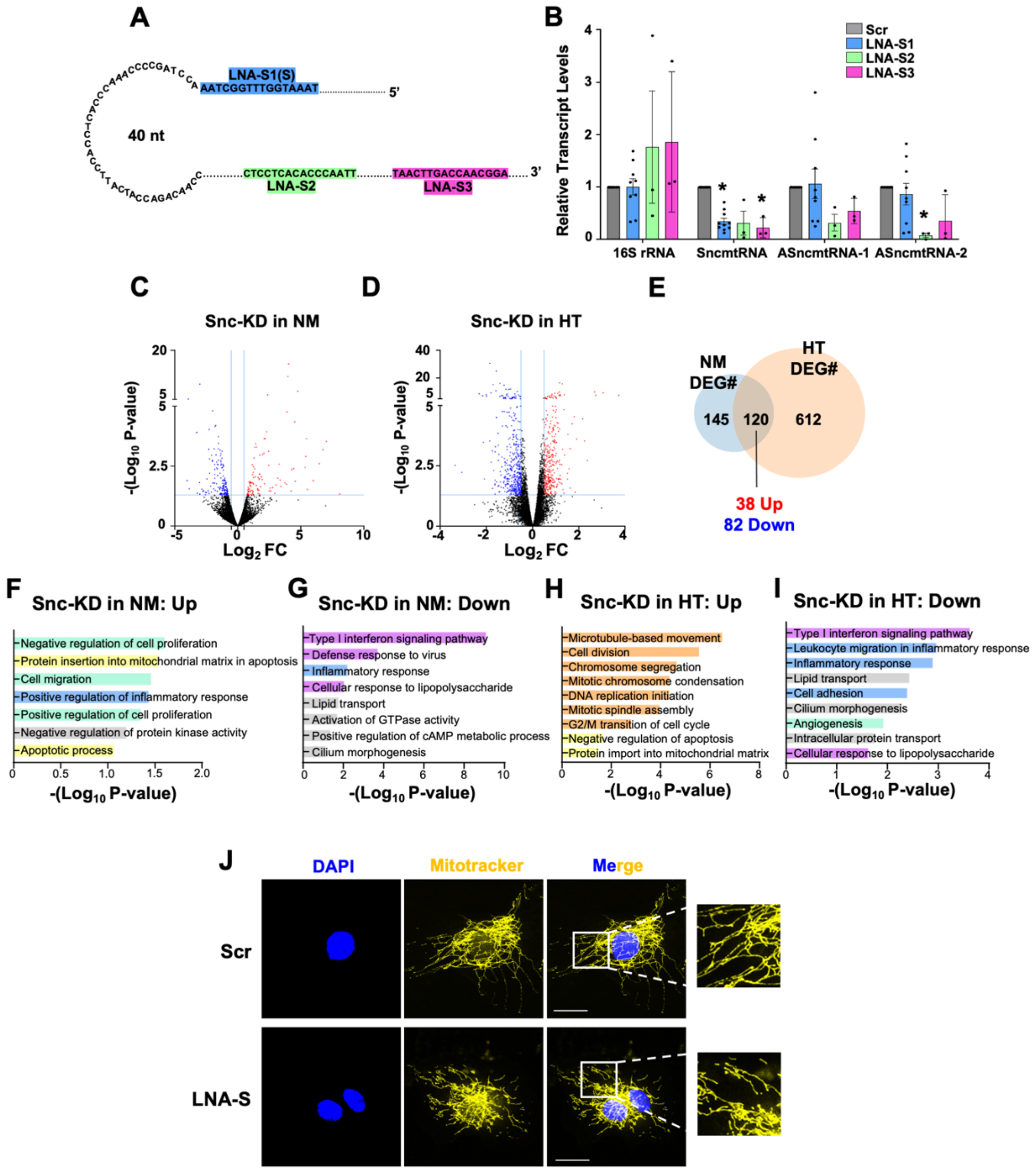


**Effect of SncmtRNA KD. (A)** Illustration of regions in SncmtRNA targeted by LNA-S1 (i.e. LNA-S), S2, and S3. **(B)** RT-qPCR of *MT-RNR2* derived transcripts in ECs transfected with respective LNA GapmeRs. **(C, D)** Volcano plots showing the DEGs ($P < 0.05$ and $Log_2FoldChange > |0.5|$) upon Snc-KD in ECs under NM (C) and HT (D). **(E)** Venn Diagram showing the number of common DEGs upon Snc-KD under NM and HT. **(F, G)** GO terms enriched from up-regulated (F) and down-regulated DEGs (G) upon Snc-KD in NM. **(H, I)** GO terms enriched in up-regulated (H) and down-regulated DEGs (I) upon Snc-KD in HT. **(J)** Representative images of Mitotracker deep red staining (yellow pseudocolor) in LNA GapmeR-transfected ECs from two independent experiments. Scale bar = 20 μm.

Fig. S3.

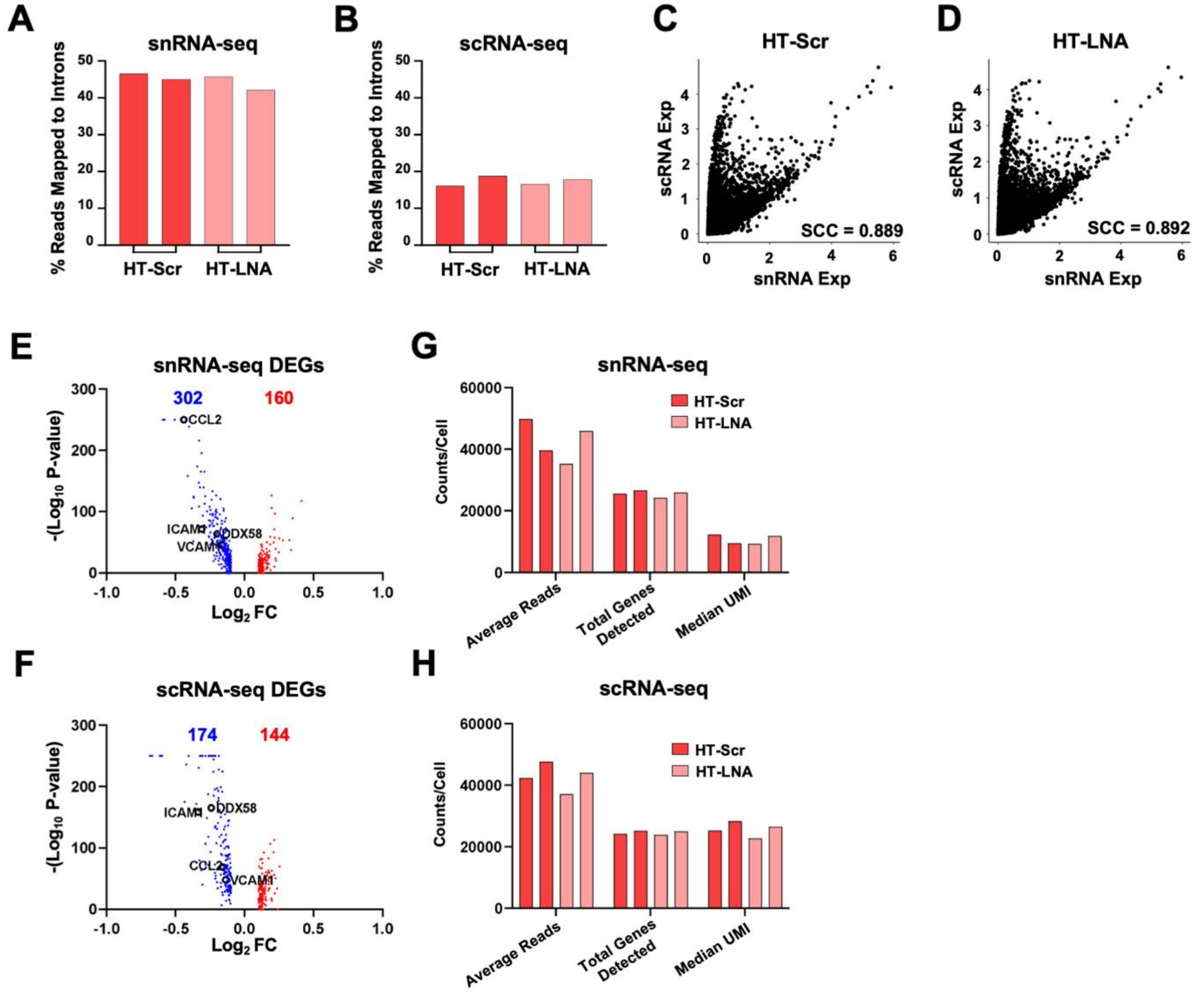


**Read mapping, correlation, and number of DEGs of snRNA- and scRNA-seq. (A, B)** Percentage of reads confidently mapped to intronic regions in snRNA-seq (A) and scRNA-seq (B). **(C, D)** Correlation plot between expression (exp) of all genes commonly captured by sn- and sc-RNA-seq in HT-treated HUVECs without or without SncmtRNA-KD, i.e., HT-Scr (C) and HT-LNA(-S) (D). SCC = Spearman correlation coefficient. **(E, F)** Volcano plots showing DEGs from snRNA-seq (E) and scRNA-seq (F). Red denotes up-regulated and blue denotes down-regulated DEGs. **(G, H)** Average reads per cell, total genes detected, and the median UMI per cell from snRNA-seq (G) and scRNA-seq (H).

**Fig. S4**

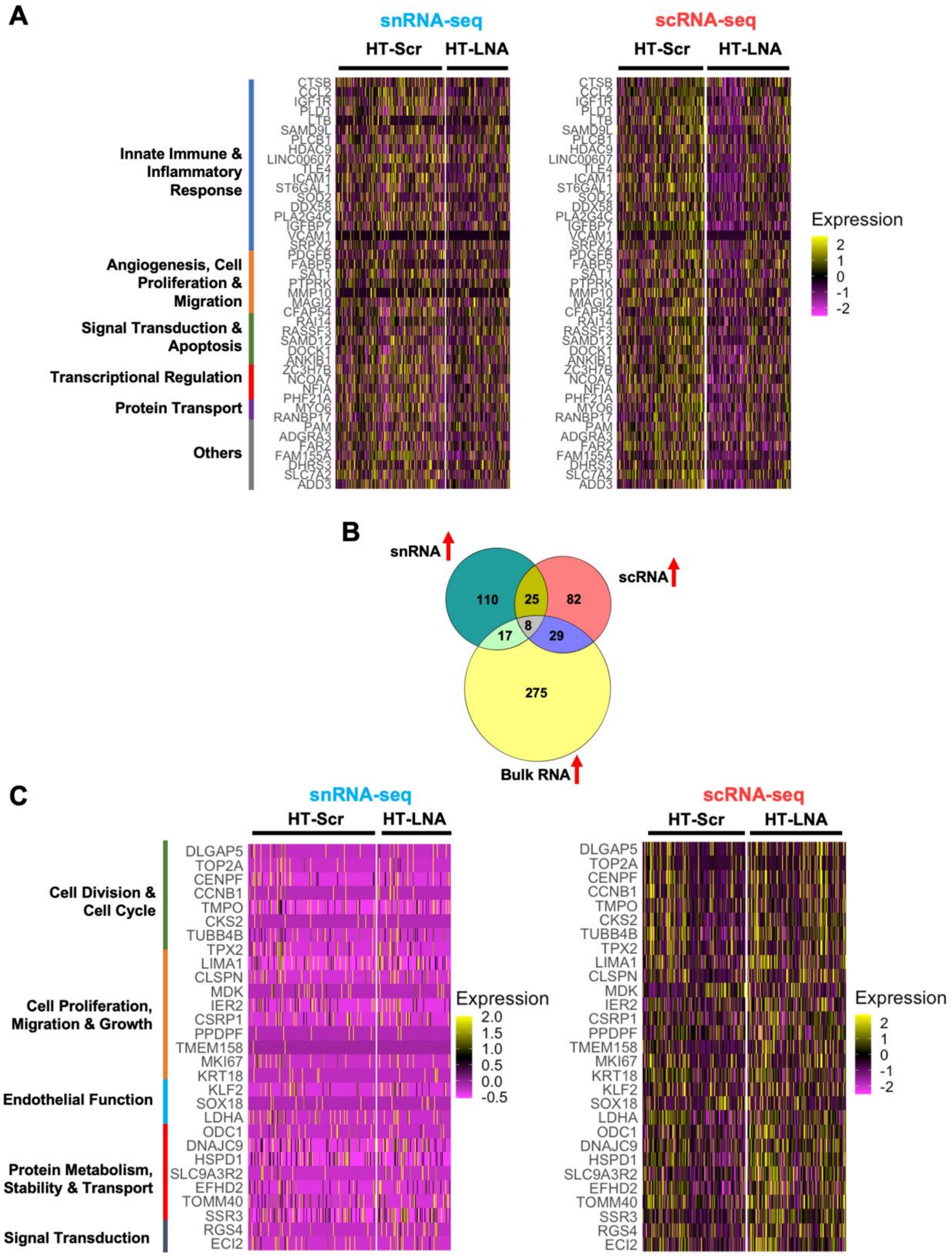


**DEG analysis of snRNA- and scRNA-seq data.** (**A**) Heatmaps showing the expression of 43 DEGs commonly down-regulated by Snc-KD in scRNA and snRNA-seq in ECs under HT. (**B**) Venn diagram showing the intersection of DEGs up-regulated in bulk RNA-, scRNA-, and snRNA-seq by Snc-KD in ECs under HT. (**C**) Heatmaps showing the expression of 29 genes commonly up-regulated in bulk RNA- and scRNA-seq but not in snRNA-seq by Snc-KD in ECs under HT.

**Fig. S5.**

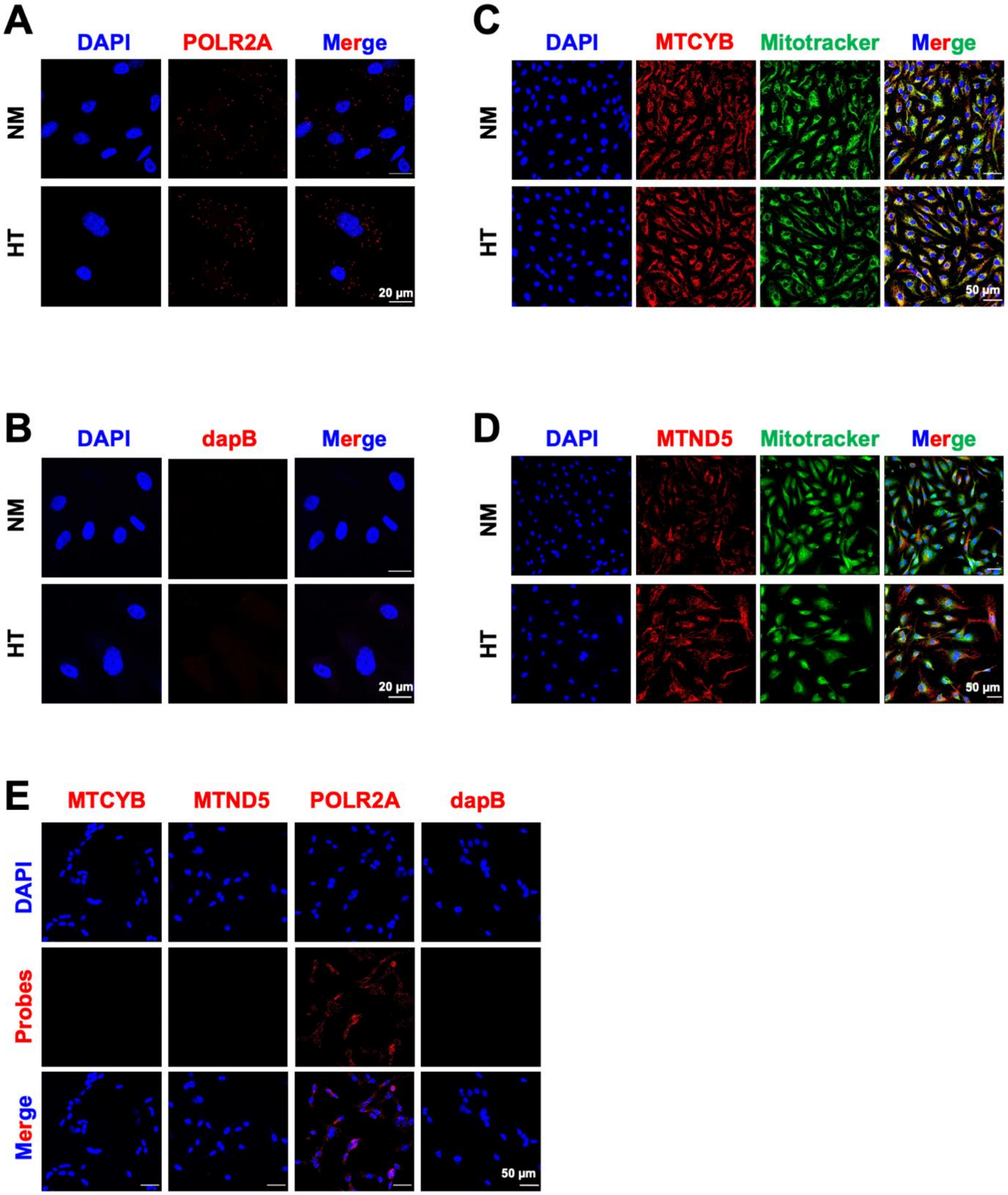


**Control experiments of smFISH for MT-CYB and MT-ND5.** (**A, B**) Confocal images of *POLR2A* mRNA (in red) as positive control (**A**) and bacterial *dapB* mRNA (in red) as negative control (**B**). (**C, D**) Confocal images of Mitotracker Deep Red (in green pseudocolor) and smFISH for MTCYB (in red) (**C**) and MTND5 (in red) (**D**) transcripts. (**E**) smFISH performed on $Rho^0$ cells for indicated transcripts, with DAPI staining the nuclei.

**Fig. S6.**

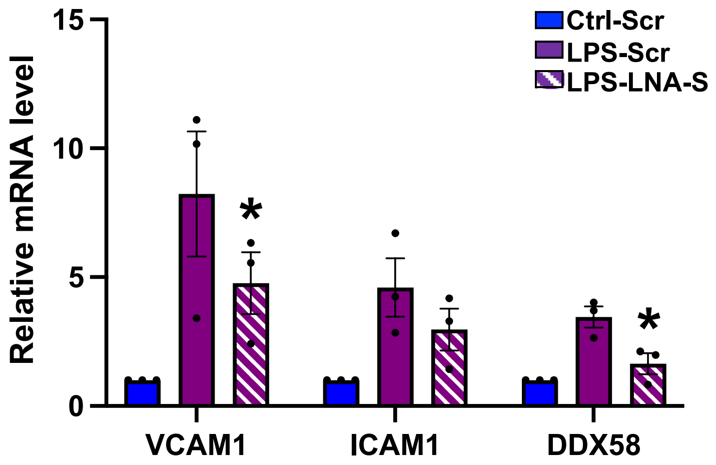


**Effect of SncmtRNA-KD in LPS-induced genes.** HUVECs were transfected with Scr or LNA-S targeting SncmtRNA and treated with LPS at 100 ng/ml for 24hrs. mRNA levels of indicated transcripts were quantified, data represent mean ± SEM from three independent experiments. *P<0.05 compared to LPS-Scr.

**Table S1. Sequences of primers, LNA GapmeRs, and FISH probes**

| Primers | Forward | Reverse |
|---|---|---|
| SncmtRNA | GGGGGTCTTAGCTTTGGCTCTCC | AAGGTGGAGTGGGTTTGGGGCTAGG |
| 16S rRNA | TTTGCAAGGAGAGCCAAAGC | ATTTAGAGGGTTCTGTGGGC |
| b-Actin | CATGTACGTTGCTATCCAGGC | CTCCTTAATGTCACGCACGAT |
| ICAM1 | GTGTCCTGTATGGCCCCCGACT | ACCTTGCGGGTGACCTCCCC |
| VCAM1 | GTCAATGTTGCCCCCAGAGA | TTTTCGGAGCAGGAAAGCCC |
| DDX58 | TGTGCTCCTACAGGTTGTGGA | CACTGGGATCTGATTCGCAAAA |
| eNOS | TGATGGCGAAGCGAGTGAAG | ACTCATCCATACACAGGACCC |
| MALAT1 | GTGCTACACAGAAGTGGATTC | CCTCAGTCCTAGCTTCATCA |
| TUG1 | CTGTGACCCAGAAGAGTTAAG | CATATCCCAGGGACTCAAAC |
| GAPDH | CTCCTCACAGTTGCCATGTA | GTTGAGCACAGGGTACTTTATTG |

| LNA GapmeRs | Sequence |
|---|---|
| SncmtRNA (LNA-S1) | TTAGCCAAACCATTTA |
| SncmtRNA (LNA-S2) | AATTGGGTGTGAGGAG |
| SncmtRNA (LNA-S3) | TCCGTTGGTCAAGTTA |
| Scramble | AACACGTCTATACGC |
